# Labelizer: systematic selection of protein residues for covalent fluorophore labeling

**DOI:** 10.1101/2023.06.12.544586

**Authors:** Christian Gebhardt, Pascal Bawidamann, Anna-Katharina Spring, Robin Schenk, Konstantin Schütze, Gabriel G. Moya Muñoz, Nicolas D. Wendler, Douglas A. Griffith, Jan Lipfert, Thorben Cordes

## Abstract

An essential requirement for the use of fluorescent dyes in biomedicine, molecular biology, biochemistry, biophysics and optical imaging is their (covalent) attachment to biomolecules. There is, however, no systematic and automated approach for the selection of suitable labeling sites in macromolecules, which is particular problematic for proteins. Here, we present a general and quantitative strategy to identify optimal residues for protein labeling using a naïve Bayes classifier. Based on a literature search and bioinformatics analysis of >100 proteins with ~400 successfully labeled residues, we identified four parameters, which we combined into a labeling score to rank residues for their suitability as a label-site. The utility of our approach for the systematic selection of single residues and of residue pairs for FRET experiments is supported by data from the literature and by new experiments on different proteins. To make the method available to a large community of researchers, we developed a python package called “labelizer”, that performs an analysis of a pdb-structure (or structural models), label score calculation, and FRET assay scoring. We further provide a webserver (https://labelizer.bio.lmu.de/) to conveniently apply our approach and to build up a central open-access database of (non-)successfully labeled protein residues to continuously improve and refine the labelizer approach.

## Introduction

Microscopy and spectroscopy techniques are ubiquitously used in the life sciences, in biophysical and medical assays, to investigate structure, interactions, and dynamics of macromolecules and their complexes down to the single-molecule level^[1–5]^. Many applications require specific labeling of the biomolecule of interest with fluorescent probes^[6–12]^. Whereas fluorescent proteins are the first choice for imaging applications in live-cells^[13–15]^, synthetic organic fluorophores (dyes) are often used for high sensitivity applications including single-molecule detection^[16–18]^ and super-resolution microscopy^[19–21]^. A common strategy for the (covalent) attachment of functional probes to proteins, including dyes, EPR spin probes, nanoparticles, and reactive surfaces, is via reactive linker moieties^[6,22]^.

A range of labeling strategies exists that exploit reactive groups, each with unique (dis)advantages. Coupling to amino groups in lysine residues can be achieved via N-hydroxysuccinimide (NHS)-esters, but this approach lacks specificity because of the abundance of lysine residues in proteins^[22]^. Alternatively, a terminally located His-tag or the N-terminus of the protein itself can be used for selective attachment of functional probes, with the disadvantage that the choice of labeling position is greatly curtailed^[22]^. In contrast, peptide tags (e.g., CLIP, SNAP, Halo, etc.) can facilitate covalent or enzymatic probe attachment (AP-BirA, LPXTG-SortaseA, etc.) at any desired location, but the size of tags limits applications and can impact protein function^[23]^. The most widely used strategy for site-specific labeling of proteins is, therefore, to introduce non-native cysteine residues and to label their sulfhydryl-moiety via a maleimide-conjugate of the selected probe^[22,24]^. Cysteine residues can be labeled with minimal effects on protein structure and function. Alternatively, unnatural amino acids (UAAs) can be introduced as targets for labeling. UAAs have proven particularly useful in cases where the removal of native cysteines is not possible due to their relevance (or abundance) and for live-cell labeling, where too many different proteins with cysteine residues are present^[25–30]^.

The introduction of cysteine residues or UAAs have become the methods of choice for many spectroscopic and microscopic studies of proteins, including the characterization of structural and functional dynamics by single-molecule Förster resonance energy transfer (smFRET)^[28,31,32]^ or pulsed electron-electron double resonance spectroscopy (PELDOR or DEER)^[33–36]^. Therefore, the ability to select optimal labeling sites for the introduction of suitable probes has grown in importance ^[37–39]^. Currently, labeling sites are typically selected based on manual inspection of the protein structure in a lengthy trial and error process to identify labeling sites via physicochemical intuition that are not essential for protein structure or function^[40–49]^, but that are also compatible with the assay requirements, e.g., for FRET to result in an inter-fluorophore distance close to the Förster Radius *R*_0_^[28,31,32]^. Frequently encountered problems when selecting a labeling site for fluorescent dyes (Figure 1A) range from (i) influence of the fluorophore on protein properties including altered biochemical function (Figure 1A, “Protein Properties”), (ii) low labeling efficiency (Figure 1A, “Sample Yield”), or (iii) unwanted dye-protein interactions (Figure 1A, “Spatial Orientation”), to (iv) unpredictable or unfavorable photophysical properties of the dyes at the chosen site (Figure 1A, “Fluorescence Properties”). Suitable residues for labeling must not only enable specific and efficient attachment of fluorophores, but also avoid the problems summarized in Figure 1A. Currently, the selection of labeling sites is often based on sensible rules of thumb^[50]^ selecting those residues that satisfy assay requirements (e.g., distance constraints for FRET^[52,53]^), but that are also solvent accessible^[54]^, show low conservation scores^[28]^ and are not related to protein function or the presence of fluorescence quenchers such as tryptophans^[50–52]^.

**Figure 1.**
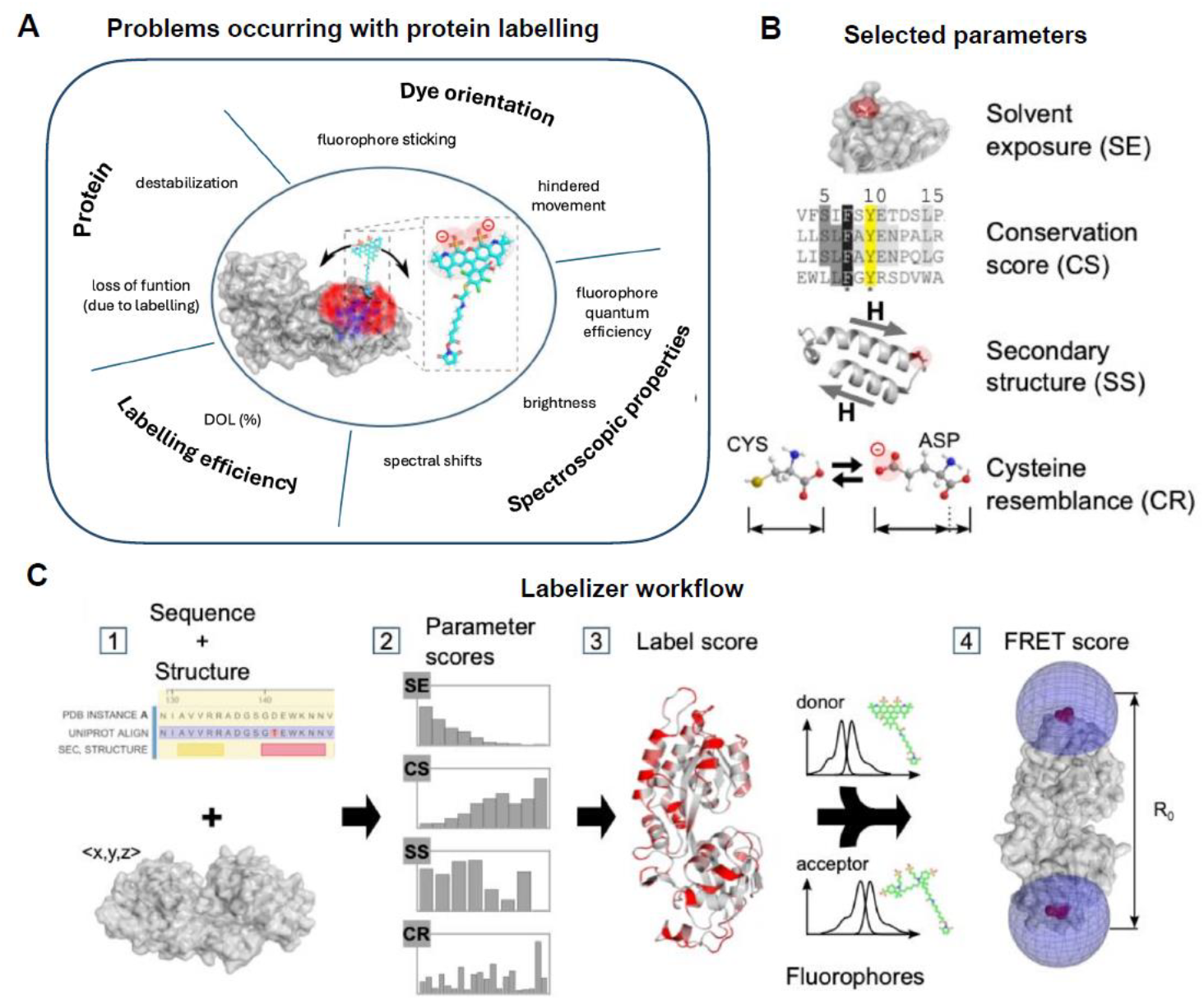
Labelizer workflow to score protein residues for labeling and FRET experiments. **A)** Schematic overview of protein-fluorophore interactions that can impact the quality and success of fluorescence assays. **B)** Parameter categories obtained from protein strctures and databases used for the labelizer analysis. **C)** Workflow for identifying suitable labeling sites, label score representation and selecting residue pairs for FRET experiments with FRET scoring.

Here, we introduce an automated analysis pipeline based on a naïve Bayes classifier^[55,56]^ to select suitable label sites using information of protein structure and sequence, e.g., from the protein data bank, PDB (Figure 1C, step 1). To systematically compare sites, we introduce a quantitative *label score LS*, which indicates the suitability of a protein residue to become a label-site, at which any of the problems shown in Figure 1A are minimal. We assembled a database of publications that report successful labeling of protein variants used in biophysical assays and identified an ideal set of parameters to allow ranking of such residues (Figure 1C, step 2/3). *LS* can be calculated independently of the choice of label (fluorophore, EPR probe, beads, surfaces etc.), yet we here focus on the use and characterization of *LS* for the attachment of fluorescent dyes to proteins. We also extended our analysis to pairs of residues for FRET assays, where the interdye distance should be close to the Förster radius to obtain highest sensitivity (Figure 1C, step 4). Therefore, we score different residue pairs according to *LS* and simulated distances to obtain an optimal FRET assay, which express the suitability of a residue pair as a FRET score. We support the predictive power of the *LS* and FRET scores with data from the literature and experiments on substrate-binding proteins (SBPs)^[57–59]^.

To make the analysis routine available to a large community of researchers, we introduce a python package called “labelizer”, which implements our analysis of protein structures, label score calculation, and FRET assay scoring. The labelizer package allows researchers to build on our findings and adapt the code for their specific needs. For straightforward use, we also provide a webserver (https://labelizer.bio.lmu.de/) with a user-friendly interface to apply our analysis approach without any programming efforts.

## Results

### Database of successfully labeled residues

As the basis of our label-site selection tool, we created a database of proteins that have been successfully labeled with fluorophores. A large set (>1000) of peer-reviewed papers and preprints was screened for labeled cysteine or UAA residues in proteins. We include protein residues in the database that have been covalently and site-specifically labeled at cysteines (predominantly) or UAAs with organic fluorophores^1^. Furthermore, only residues are included for which the structure of the protein has been deposited in the PDB. For the included proteins, we extract information on the labeled residue (chain, number), the type of mutation used for labeling (cysteine or UAA), the assay type (e.g., single fluorophore assays, smFRET assay with two labels, imaging, bulk FRET, etc.), and the type of label. We then gathered additional information on the protein, such as its oligomeric state (monomer, dimer, complexes), whether the protein structure has been experimentally determined or only a homology model is available, and whether it is a soluble or a membrane protein. Overall, we identified labeled residues in >100 different proteins from >100 publications (see Supplementary Data: Reference Database Labelizer). An overview of the data and summary statistics are presented in Supplementary Figure S1.

We used a standardized pre-processing routine (see Methods and Supplementary Note 1) to extract all relevant residues from the pdb-files of the proteins in the database. The final data set from 104 pdb structures contains 43357 residues, 396 of which are reported to have been successfully labeled (the other residues are considered unknown). For all residues in our database, we compute multiple parameters that can be assigned to one of the four major categories (Figure 1B): (i) conservation score CS (ii) solvent exposure SE, (iii) secondary structure SS, and (iv), amino acid similarity of the exchanged residues to a cysteine, which we abbreviate as cysteine resemblance CR (see Supplementary Note 1 with Tables S1-S6). The parameters are either directly extracted from the residues in question, e.g., amino acid type, mass, charge and size, or calculated with the help of freely available software (conservation score (ConSurf^[60,61]^), solvent exposure (DSSP^[62]^, HSE^[63]^, MSMS^[64]^), and secondary structure (DSSP^[62]^)). Altogether, we obtain 28 parameters for each residue.

### Bayesian approach to the prediction of labeling sites

To identify suitable residues for labeling, we are interested in *P*(*l*|*s*), the conditional probability that the residue can be labeled given a parameter value *s*. By Bayes’ law

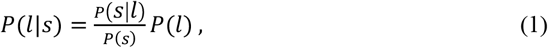

*P*(*s*) is the probability distribution of the parameter values *s* over all residues, whether or not they can be labeled, while *P*(*s*|*l*) is the probability distribution of the parameter values *s* given that the residue can be labeled. Finally, *P*(*l*) is the *a priori* probability that a residue can be labeled. While *P*(*s*) and *P*(*s*|*l*) can be readily computed from our database of labeled protein structures, *P*(*l*) is harder to assess, since the literature is biased towards reporting successful attempts of labeling that have provided relevant insights. Since *P*(*l*) only scales the final probability and does not affect the predictions of the relative ease of labeling for different residues, we decided to here use a simplified parameter score

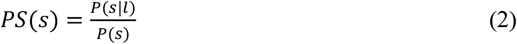

instead of *P*(*l*|*s*) to assess the suitability of residues for labeling. *PS(s)* is in essence the odds ratio for a given parameter value to occur in a labeled residue compared to randomly selected residues. For all 28 parameters, we computed *P*(*s*|*l*) distributions for the 396 successfully labeled residues and *P*(*s*) distributions from all 43357 residues of the 112 chains of the database (Figure 2A and Supplementary Figure S2/3).

**Figure 2.**
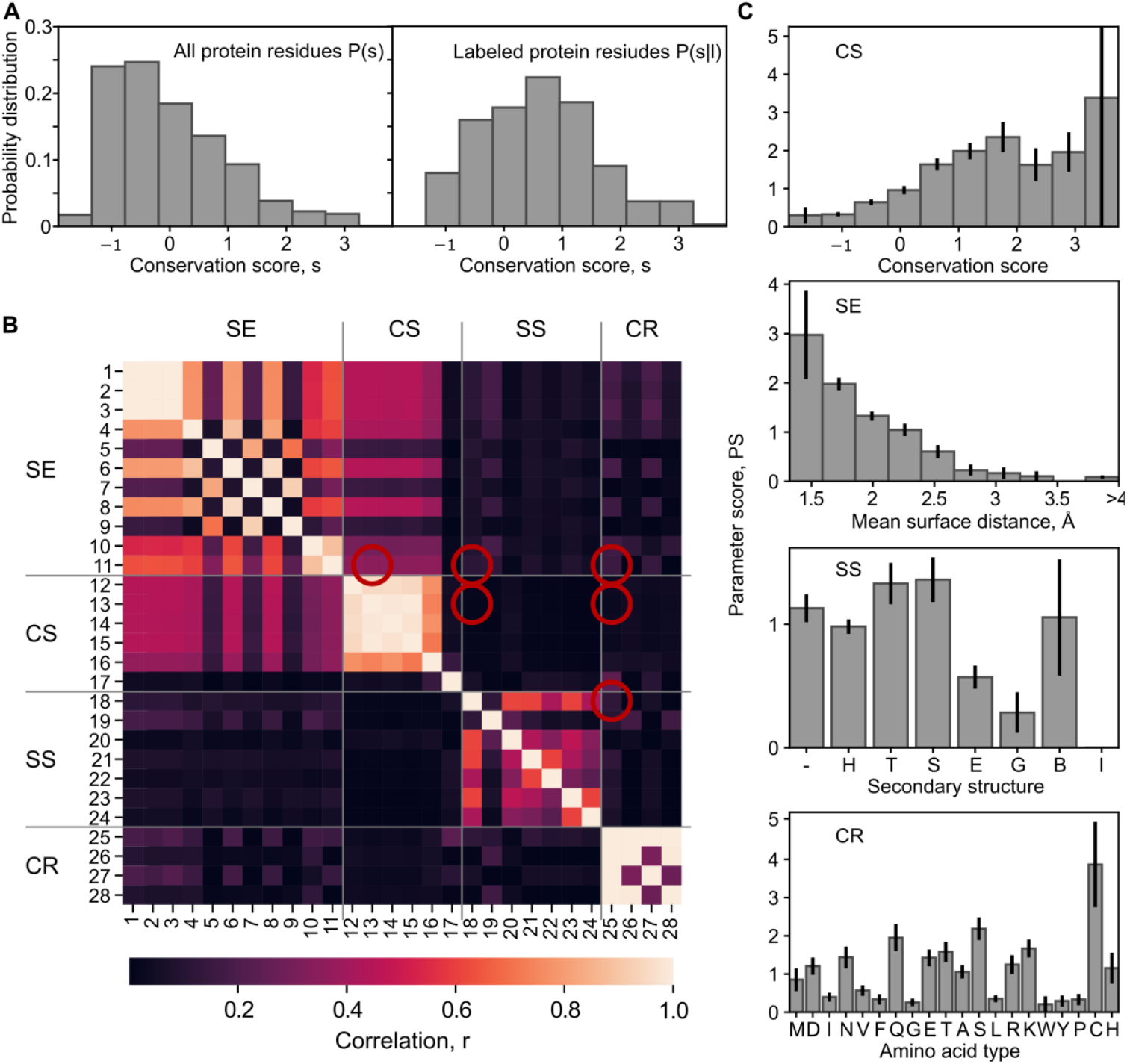
Parameter score analysis. **A)** Probability distribution *P*(*s*) for the parameter ConSurf score (#13, Supplementary Table S2 and S6, negative values represent highly conserved residues among homologues) for all analyzed residues (left) and for the successfully labeled residues *P*(*s*|*l*); right. **B)** Correlations between all parameters were calculated based on Pearson correlation (numeric-numeric), interclass correlation (categorical-numeric) or Cramer’s V (categorical-categorical). The cross-correlations of the final parameter selection for the labelizer algorithm are marked (red circles). **C)** Parameter score distributions *PS* = *P*(*s*|*l*)/P(*s*) for the four parameters that we select as the default for scoring. The top panel shows the resulting parameter score from the distribution in A. For the other categories, the parameters are: solvent exposure (#11, mean surface distance, Supplementary Table S1/S6), secondary structure (#18, secondary structure from DSSP, Supplementary Table S3/S6), and cysteine resemblance (#25, amino acid identity, Supplementary Table S4/S6). Error bars are the standard deviation from counting statistics. The clear deviations from uniform distributions indicate that all four parameters contain information about the suitability of a site for labeling.

As a control, we compared the probability distributions *P*(*s*) from our database of successfully labeled residues with the distributions computed for a random selection of protein chains from the PDB (PDBselect, November 2017)^[65,66]^ (see Methods). Here, we find only minor differences, indicating that the protein parameters in our database are representative of the overall PDB content (Supplementary Figure S2). One notable difference is that cysteines are much less abundant (by ~50%) in the database of labeled proteins compared to the overall PDB, suggesting that cysteine insertion and labeling is easier (or at least more common) for proteins with fewer native cysteines (Supplementary Figure S2). Although we also included residues that were labeled via UAA incorporation, our database indicates that cysteine labeling is still the predominant strategy for proteins, since it makes up ~90% of all labeled residues in our database (Supplementary Figure S1D).

We find clear differences between *P*(*s*|*l*) and *P*(*s*) and, therefore, non-uniform *PS* distributions for most of the investigated parameters (Figure 2A/C and Supplementary Figure S6), showing that they indeed contain information about the suitability of residues to serve as label sites. To evaluate which parameters are most predictive, we computed *PS* distributions for 28 parameters (numbered from #1 to #28) from all four categories from our database (Figure 2 and Supplementary Table S1-4). For each *PS* distribution, we analyzed their mean-square deviation from an equal distribution, the Gini coefficient, and the Shannon entropy (see Supplementary Note 1 and Supplementary Table S6). We find that the *PS* distributions for many parameters clearly deviate from an equal distribution and contain significant information (low Shannon entropy), e.g., seen in #1: relative surface area (Wilke), #4: first half-sphere exposure (10 Å), #16: variant length in homologues (see Supplementary Figure S3). Other parameters contain barely any information such as #17 cysteine in homologues (yes/no), or #27 amino acid charge (Supplementary Figure S3). Thus, strikingly, it is largely unpredictive for labeling of a residue whether a cysteine is found in one of the homologue proteins at the same position or whether the residue is charged (see parameter #17 and #27, Supplementary Figure S3). One might have expected that residues with cysteine homologues are easily mutated to cysteines, and therefore, significantly enhanced in our scoring, which is not the case.

After establishing the predictive power of individual parameters, we investigated what combinations of parameters should be used. For this we calculated the correlation between all parameters^2^ to judge their statistical independence, which is desirable for our Baysian analysis (Figure 2B). We formed sets of four parameters and used a correlation measure (2-norm of all paired correlations, see Methods) to calculate a combined correlation estimator for all combinations of parameters (Supplementary Figure S4). Whereas this combined correlation-derived measure shows higher values for most combinations of two or more parameters within the same categories CS, SE, CR, and SS, the correlation of combinations of parameters from different categories was smaller (<0.5). This effect was independent of whether parameters with high or low predictive power (MSD / Shannon entropy) were combined (Figure 2B and Supplementary Figure S4). The overall low correlation between parameters from different categories justifies our categorization and their consideration as independent variables if we restrict our selection to one parameter per category. The strong correlation within categories also suggests that the choice of the particular parameter from one category is not critical, i.e., most of the parameters can account for the properties of the respective category.

### The combined label score predicts potential labeling sites

To combine parameter scores into a final assessment of a given residue to serve as label site, we introduce a combined label score, *LS*. By standard probability theory different parameters *s*_*i*_ can be combined by

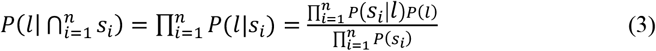

under the assumption that they are independent, where Π denotes the product and ∩ the intersection. This naïve Bayes classification^[55,56]^ is known to give good predictions for low and moderately correlated parameters^[67–70]^, which is the case for our parameter set (Figure 2B). In general, any residual correlation alters the calculated probability values towards the extremes of 0 and 1^[70]^. However, we again use parameter scores as comparative figures without the meaning of probabilities and combine the *PS*_*i*_ into the combined label score by taking their geometric mean:

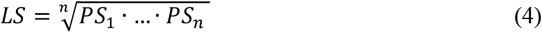

An important question is which of the 28 parameters to include in the LS. We include one parameter from each of the four categories CS, SE, SR, and SS, for which concrete values were mapped onto the structure of the phosphate binding protein PBP (Figure 3A). For a rational selection of parameters, we strive (i) to maximize the dynamic range of values for *LS*, (ii) to maximize the enhancement/suppression level of *LS* of the successfully labeled residues in the database for high/low *LS* values and (iii) to maximize the statistical significance level of *LS* values of random residues over *LS* values of the labeled residues in the database.

**Figure 3.**
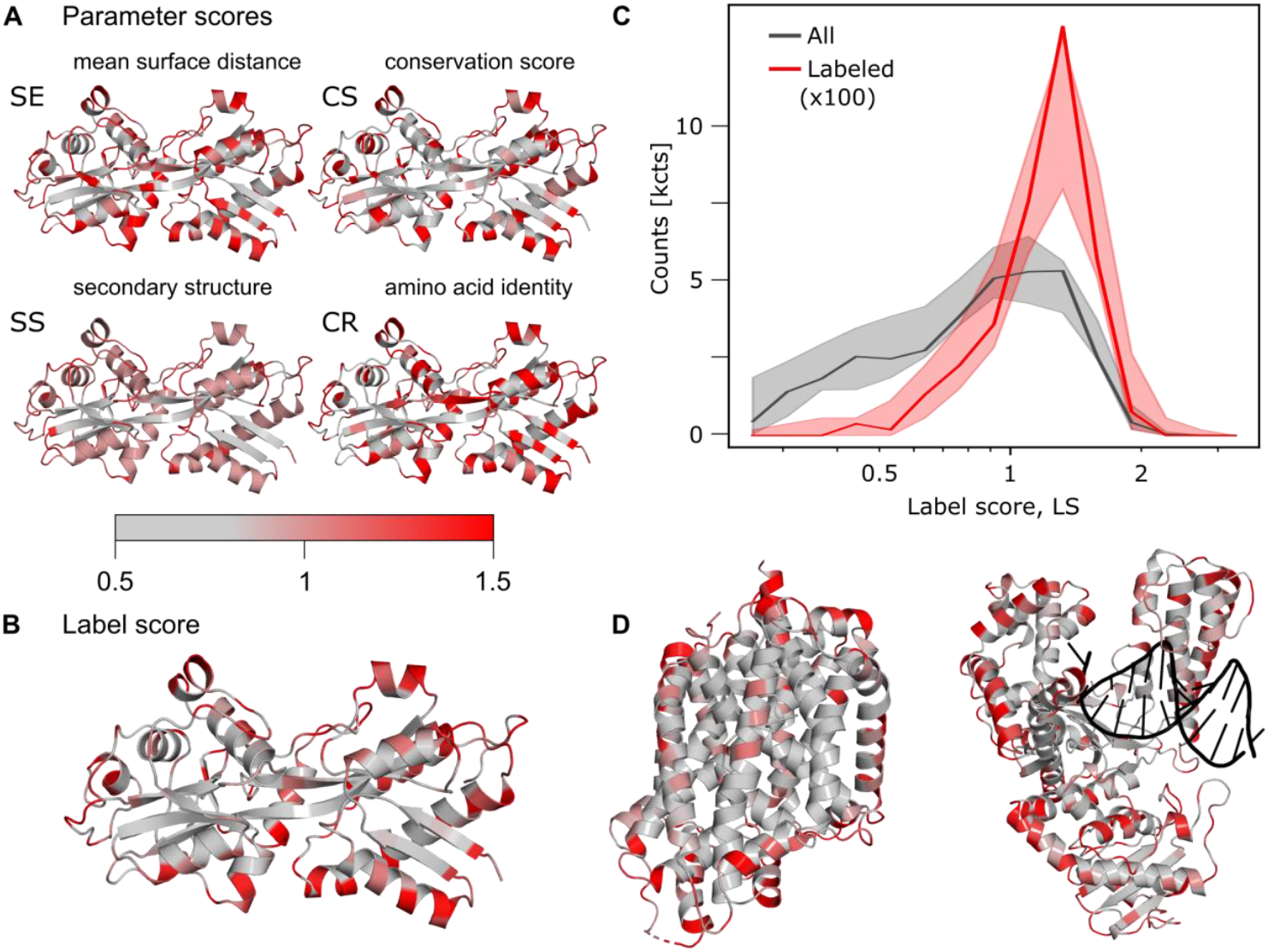
Visualization of parameter and label scores. **A)** Visualization of the selected parameter scores from the four categories, which are used as default settings in our webserver for the example of PBP from *E*.*coli*^[71,72]^; pdb:1OIB. **B)** Visualization of the label score on PBP based on default parameters shown in panel A. **C)** Label score histogram of all residues (gray) and labeled residues (red) in the database. Due to lower numbers of residues in the labelled data set it was multiplied with a factor of 100 to allow better comparison of the distributions. The shaded area shows the 95% confidence interval from 400 bootstrapping runs. **D)** Additional examples of *LS* values indicated on protein structures for a membrane protein (left, LeuT of in A. aeolicus^[73,74]^, pdb:2A65) and a DNA-binding protein (right, DNA polymerase I of B. stearothermophilus^[75,76]^ with DNA template, pdb:1L3U).

Based on these criteria, we were able to identify several parameter sets with predictive power (Supplementary Figure S5A,B), but also combinations with much less information (Supplementary Figure S5C). In the end, we decided on one set that resulted in a large difference of the distributions between the random and labeled residues: mean surface distance (SE, #11), conservation score (CS, #13), secondary structure of the labeled residue (SS, #18), and the mutated amino acid (CR, #25). This set is shown in Figure 3C and is used as default for *LS* calculations in this manuscript and for the associated webserver. In the labelizer python package any parameter combination can be selected.

We chose the default set out of all well-performing combinations, because of the intuitive nature of all selected parameters and the maximized differences between the mean *LS* values of all vs. the labeled residues. Both our choice of parameters and the selected number of categories to four (and not only two or three) are supported by statistical analysis of the significance, i.e., a *t*-test and a comparison of the mean values of all vs. labeled parameters for different parameter combinations (Supplementary Table S7). Our selection is further validated by comparing the receiver operating characteristic (ROC curve) for the baseline, when retraining with one of four scores removed and the predictive power of each of the scores on their own (Supplementary Figure S6). Bootstrapping of the final set demonstrates the robustness of our analysis (Figure 3C). For this final set of parameters, we find that the label scores *LS* range from 0.2 to 2 for most residues (except 5% failed calculations with *LS* = 0). The ratio of the *LS* distribution of successfully labeled residues in the database and all label scores shows that high label scores (>1.5) are significantly enhanced by a factor of ~3-4 for the labeled residues, whereas low label scores (<0.5) are suppressed by a factor of ~10 (Figure 3C). This suggests that the label score is an informative measure to rank and compare residues for their suitability for labeling with fluorophores.^3^ We visualize the calculated *LS* scores for three typical proteins, comprising a soluble protein, a membrane protein, and a DNA binding protein (Figure 3D-F).

### Experimental benchmarking of the label score

To characterize the relation between *LS* values and experimentally observed behavior, we performed two different analyses of variants of the maltose binding protein (MalE) with single-cysteine labeling sites. MalE is a soluble bilobed protein with an open (apo) and a closed (holo) structure^[59,77]^, which serves as the periplasmic component of the bacterial ABC importer MalFGK_2_-E^[78]^. We visualized *LS* values for all sites of apo MalE in Figure 4A and the corresponding distribution in Figure 4B. The distribution shows a *LS* value range between 0 and 2; high values for *LS* appear mostly in positions of MalE near the surface, when *LS* are mapped back to the structure (Figure 4A).

**Figure 4.**
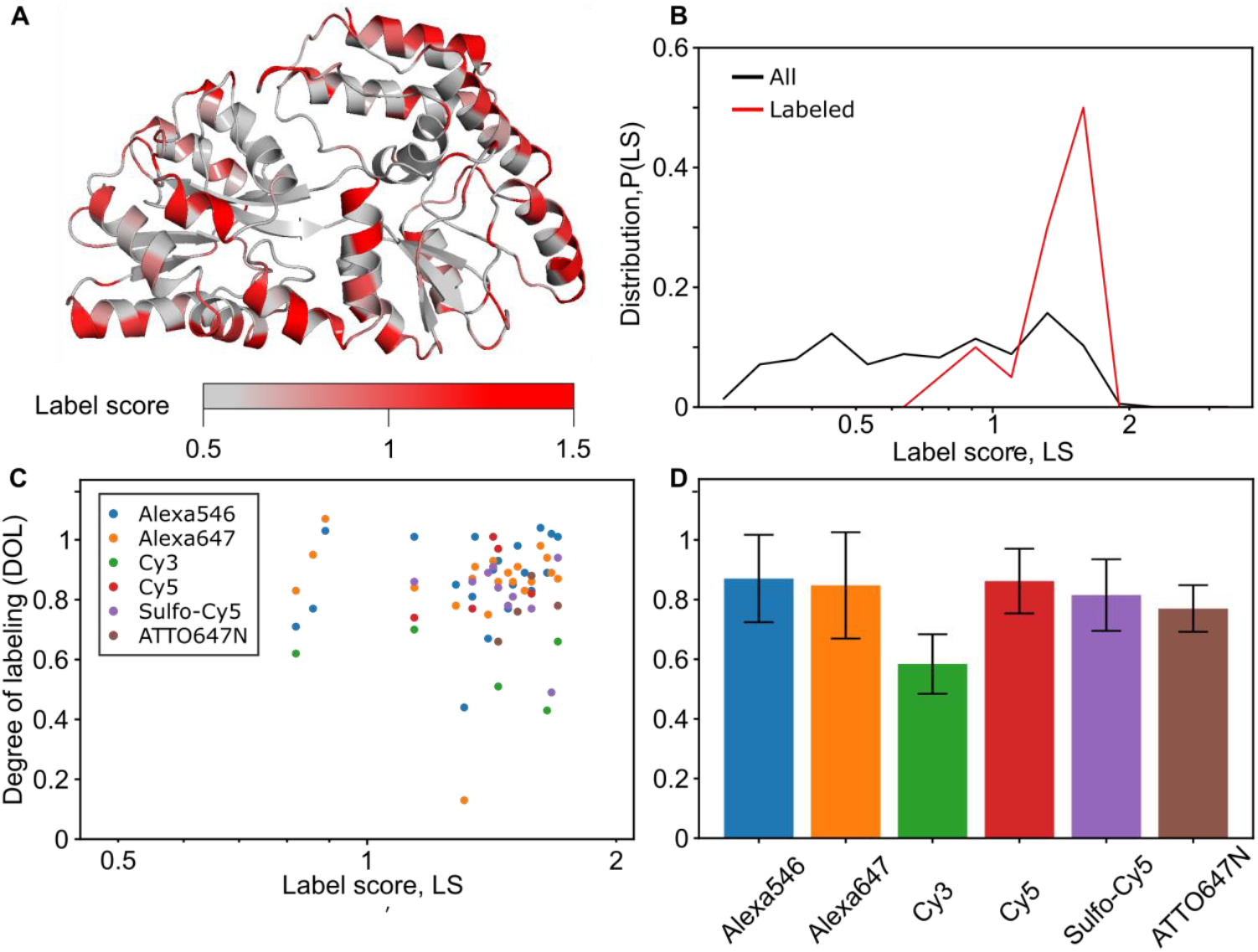
Characterization of LS values with experimental parameters and degree of labeling (DOL) for single cysteine variants of MalE. **A)** Crystal structure of MalE in the apo state with LS color coded. **B)** LS distribution of apo MalE for all (grey) and the 20 successfully labeled residues (red). **C)** LS vs. DOL for the MalE labeling data set, where different dyes are color coded. **D)** Average DOL for different dyes, based on the data in panel C. Error bars indicate the standard deviation. We do not observe a correlation of label efficiency and label score for the selected mutants with in general high label scores LS>1. We do not observe a significant difference across fluorophore types (p-value >0.05) except of Cy3, which showed significantly lower DOL values (p-value <0.05) compared to all other fluorophores.

First, we studied a data set of 20 variants of MalE, partially taken from previous work with (relatively) high LS scores representing residues that are good candidates for labelling according to our approach. For these experiments, we used the dyes Alexa546, Alexa647, Cy3, Cy5, sCy5 and ATTO647N and obtained an average degree of labeling (DOL) of 0.82 over all samples after protein labelling and SEC purification (Figure 4). The *DOL* was determined using the molar ratio between fluorophore and protein concentration from the Lambert-Beer law: *DOL* = c(fluorophore)/c(protein). All successfully labeled sites have an average of ~1.4 and almost 90% of them showed LS values >1 (Figure 4B/C). The distribution of the label scores for the successfully labeled sites is different from the distribution of all residues of the MalE protein (Figure 4B), again confirming that LS provides valuable information about the suitability of protein residues to act as label site. Our analysis shows, however, no correlation between LS and the experimentally determined DOL (Figure 4C). This is not too surprising since all tested residues have relatively high label scores and we focused on mutants with a reasonable chance of labeling and did not include measurements e.g., of buried residues with low label scores. Furthermore, we do not observe systematic differences between different dyes, suggesting that our method works robustly and is independent of the fluorophore (Figure 4D and Supplementary Data; *LS* vs. *DOL*).

In a second set of experiments, we ranked all MalE residues by their label scores and then randomly selected 5 variants each from the best 10% *LS* scores (referred to as “positive control”) and 5 residues from the worst 10% *LS* scores (“negative control”). For each of these 10 variants, we characterized the effect of the cysteine mutation in terms of the protein’s expression yield and DOL using the dye sCy5 (all data are provided in the Supplementary Data Excel file and Table 1). All positive controls, i.e. MalE variants comprising residues with high label scores, expressed with high yields (> 15 mg from a 2 L expression culture) and could be labeled with a DOL > 85%. These findings again support that residues with high LS scores can be successfully expressed and labeled, in line with the first analysis of MalE point mutations (Figure 4).

**Table 1.** Overview of expression and labelling properties of randomly selected MalE variants.

| Cysteine variant | Control | Expression | Labelling |
| --- | --- | --- | --- |
| Q72C | positive | OK | OK |
| S211C | positive | OK | OK |
| K219C | positive | OK | OK |
| E309C | positive | OK | OK |
| E322C | positive | OK | OK |
| L7C | negative | reduced | X |
| W94C | negative | no expression | n.d. |
| I116C | negative | OK | X |
| G228C | negative | OK | OK |
| W230C | negative | OK | OK |

In contrast, two of the negative control variants showed reduced expression yield (L7C with 7.8 mg) or no expression at all (W94C). Furthermore, two of the obtained four negative control variants showed DOL values <2%. Interestingly, the other two negative control variants showed good expression yields and adequate DOL values, suggesting that not all residues with low LS scores are necessarily unsuitable for labeling. Taking all variants from this set of 10 MalE variants into account, there is a statistically significant correlation between LS score and DOL (*p* = 0.03 from a two-sample *t*-test), further supporting the approach presented here.

### Extension of the LS score to FRET experiments

To test our prediction tool for the design of a biophysical assay we extend it to FRET experiments. For this we combine the label score *LS* with an additional parameter for the rational design of FRET experiments. The central idea is to select residue pairs for FRET experiments that are (i) suitable as label site based on *LS*, (ii) are separated by a distance that is close to the Förster radius of the dyes used (for maximum sensitivity) and (iii) that can detect conformational motion. Criteria i/ii are relevant to the case where one protein structure is available, and a residue pair is wanted with a distance close to the Förster radius of the dye pair. In this scenario, the researcher can use combinations of residues in different domains of the protein for maximal sensitivity. We define the FRET score *FS* of a residue pair *{i,j}* for a single protein structure as:

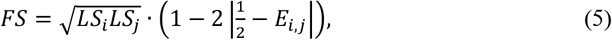

*FS* considers the label scores *LS*_i_ and *LS*_*j*_ of two residues *i* and *j* in the protein structure with corresponding predicted FRET efficiency *E*_*i,j*_ (see Supplementary Note 2 for details on the FRET efficiency prediction). *FS* is highest for residue pairs with predicted E_i,j_ = 0.5, i.e., an interdye distance similar to the Förster radius of the dye pair.

If two (interconverting) structures of a protein are available, one is interested to find FRET pairs that show the largest possible shifts in FRET efficiency. This scenario is encountered when ligand binding, protein-protein interactions or other macromolecular interactions are studied, and requires that distinct structures of the same protein, e.g., ligand-free and ligand-bound, are available. We define the FRET difference score *FS*_Δ_ of a residue pair *{i,j}* for two available structures *A* and *B* of the same protein as

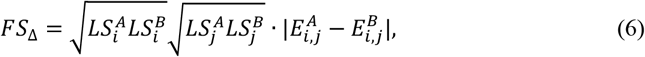

with the label scores LS of two residues *i* and *j* in two protein structures *A, B* with their corresponding FRET efficiencies 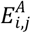 and 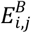, respectively.

### Accessible volume calculations for FRET labels

To rationally establish a FRET assay with maximum sensitivity it is necessary to operate at interprobe distances around the Förster radius. A crucial step for calculation of both FRET scores is, therefore, the ability to predict interdye distances from the protein structures accurately (Figure 5). The labelizer package supports three models for *in silico* fluorophore distance predictions. A rough approximation of expected FRET efficiencies can be obtained from the C_β_ distances between two residues^[54]^ (Figure 5A). However, these distances can differ >10 Å from the actual mean fluorophore positions, due to the size of the fluorophore and the flexible linkers (10-20 Å length) used for fluorophore attachment^[79,80]^. While distance changes are less impacted by such deviations, the absolute distances are significantly affected by the geometry of the labels (Figure 5C). Neglecting these effects can reduce the sensitivity of a FRET assay by up to a factor of ~4 (Figure 5D and Supplementary Figure S7 and 8).

**Figure 5.**
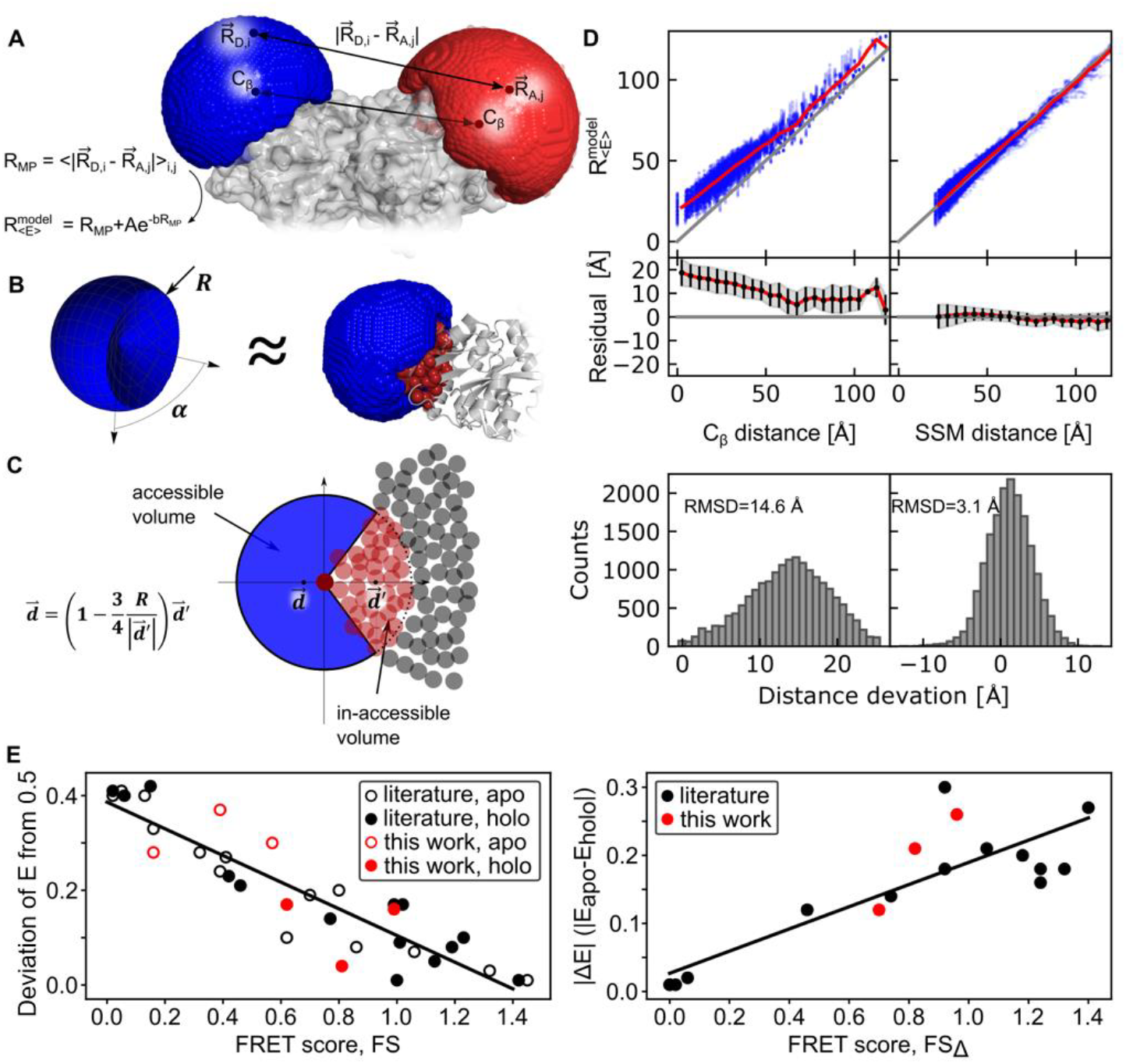
Accurate prediction of interdye distances on proteins and experimental benchmarking of the FRET scores. **A)** Distance estimation with FPS computes a grid-based accessible volume to determine the mean-position of the fluorophore 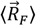, the averaged inter-fluorophore distance 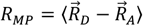, and the efficiency weighted average fluorophore distance 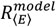 approximated with an exponential correction factor (see Supplementary Note 2). Illustration shows donor and acceptor labeling at residues S3C and P86C, respectively, in PBP (pdb: 1OIB). **B)** Approximation of the accessible volume with a spherical sector. The spherical sector (left) is defined by the radius R (linker length of the fluorophore) plus an opening angle *α* and approximates the accessible volume simulated with FPS software (right, blue volume). The red spheres represent the protein atoms within radius R from the C_ß_ atom. **C)** Illustration of the determination of the mean position in 2D. The circles represent the atoms of the protein. The inaccessible volumes are the atoms within a radius R (pale red circle) to the C_ß_ atom (dark red circle). **D)** Comparison of C_ß_ distances and modelled distances with the introduced spherical sector approximation compared to FPS-derived distances in 10 selected pdb structures with 35 different fluorophore parameters (*N* = 32116, top). The data >100 Å are noisy due to low statistics. Histogram of the distance offsets from the C_ß_ atom for the distances in the range 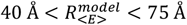 (*N* = 17359, bottom). **E)** Analysis of the deviation of measured FRET efficiencies from 0.5 for selected MalE mutants taken from ref. ^[39,81]^ (literature) and new experiments (this work) in apo (empty circle) and holo state (full circle) with respect to the computed FRET scores (left). Measured FRET efficiency shift between apo and holo of mutants in A plotted against the FRET difference score *FS*^Δ^ (right). Linear fits of the data are shown as solid lines with R^2^ = 0.84 (left) and R^2^ = 0.73 (left).

To predict distances between fluorophore labels accurately, it is important to obtain accurate simulations of the accessible volumes (AVs) considering the size and shape of the dyes and their linkers. Molecular dynamics simulations have been successfully used for this purpose^[82–84]^, yet they are too slow as a screening tool. Coarse-grained simulation via FRET-restrained positioning and screening system (FPS), where all positions on a grid are examined to decide whether it can be occupied by a fluorophore of specified size and linker length, provide AVs that are in very good agreement with experimental values of interdye distances^[49,80,81,85–90]^ (Figure 5A). Comparing the calculated C_ß_-distances of the residues with FRET-averaged distances 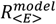 from AV simulations reveals deviations of 10 to 15 Å (RMSD, Figure 5B and Supplementary Figure S8A), highlighting the need to consider the dye and linker geometry. The computation time required for one pair of dyes using FPS, however, is still rather long for screening purposes, e.g., several hours when >10.000 residue pairs should be considered (see Supplementary Table S9).

Therefore, we here introduce a simpler and faster distance estimator based on a spherical sector model (SSM) that estimates dye-accessible and dye-inaccessible volumes (Figure 5B). SSM is used for screening purposes since it is 100 to 1000 times faster than currently available simulations such as FPS. Our algorithm relies on an approximation of the accessible volume by a spherical sector of angle *α* and radius *R* representing the linker length of the fluorophore (see Figure 5C). The atoms of the protein within radius *R* from the attachment site (C_ß_ atom) define an inaccessible volume (see Figure 5B/C, pale red spheres). We find a direct relation between the center of mass of these atoms 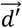 (inaccessible volume) and the center of mass of the accessible volume 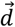 (see Supplementary Note 2) as

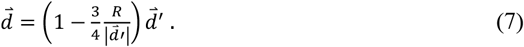

We included a small correction *ε* (~0.5 Å for typical fluorophores) to the linker length 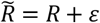 in this formula to compensate for the size of the fluorophore core (Supplementary Note 2, Supplementary Figure S7) and we used an estimation to convert the distance of the mean positions to FRET-averaged distances (Supplementary Note 2, Supplementary Figure S8). To test our method, we performed distance simulations for 100 donor-acceptor pairs in 10 different protein structures, where we altered the linker length and the dye dimension with 35 variations resulting in 35.000 distance simulations in total. Our SSM approach gives results in good agreement with the FPS method with a deviation of ±3 Å (RMSD, Figure 5 and Supplementary Figure S7), which is on the order of the intrinsic distance precision of FRET^[80]^. The mean-position distances are converted to FRET-averaged distances with an exponential correction factor at small distances (see Methods and Supplementary Figure S8). The spherical sector method allows to screen >10.000 FRET-pairs within seconds on a single CPU with <1 ms calculation time per residue-pair (see Supplementary Table S9). Therefore, our standard settings are to use the SSM method for a first selection of suitable FRET-labeling positions and subsequently refine the best three hundred FRET pairs with the FPS AV-simulations^[81,90]^. Alternatively, our python package allows calculating the C_ß_ distances (low accuracy) or the FPS-derived derived distances (long runtime) for all residues by manual selection.

### Experimental benchmarking of the FRET score

At first we used the labelizer workflow to establish new FRET assays for mechanistic studies of the ABC transporter-related prokaryotic substrate-binding protein PBP^[57–59,91]^ (Figure 6A). As seen in the crystal structures PBP undergoes a ligand-induced transition from a ligand-free open (pdb: 1OIB, apo) to a ligand-bound closed state (pdb: 1PBP, holo; Figure 6A). Yet, the ligand binding mechanism of PBP, i.e., ligand-binding before conformational change (induced fit) or conformational change before ligand binding (conformational selection) has not been studied. Thus, our goal was to obtain assays with large changes in FRET efficiency upon addition of the ligand inorganic phosphate for dye pairs with a Förster radius around 5 nm. We identified multiple suitable residue combinations with maximized positive and negative distance changes based on *FS*_Δ_ (Figure 6B). We selected four double cysteine variants with large predicted shifts from long (low FRET) to shorter (higher FRET) distances upon phosphate binding. We selected those from the list of 300 refined pairs using the FPS parameters for Alexa Fluor 555-Alexa Fluor 647 (Figure 6B; Supplementary Table S8). Before conducting FRET experiments, we characterized one of the double-cysteine variants PBP (S3C-I76G-P86C) and the cysteine-less PBP variant PBP (I76G) biochemically by ITC and obtained K_d_-values of 10 ± 5 µM for PBP (S3C-I76G-P86C) and 19 ± 6 µM for PBP (I76G); (mean ± SD from n = 2 protein preparations (Supplementary Figure S9). These experiments suggest that protein labelling does not affect substrate affinity. The I76G mutation was used in all PBP protein variants presented in this paper (Figure 6 and Supplementary Figure S9).

**Figure 6.**
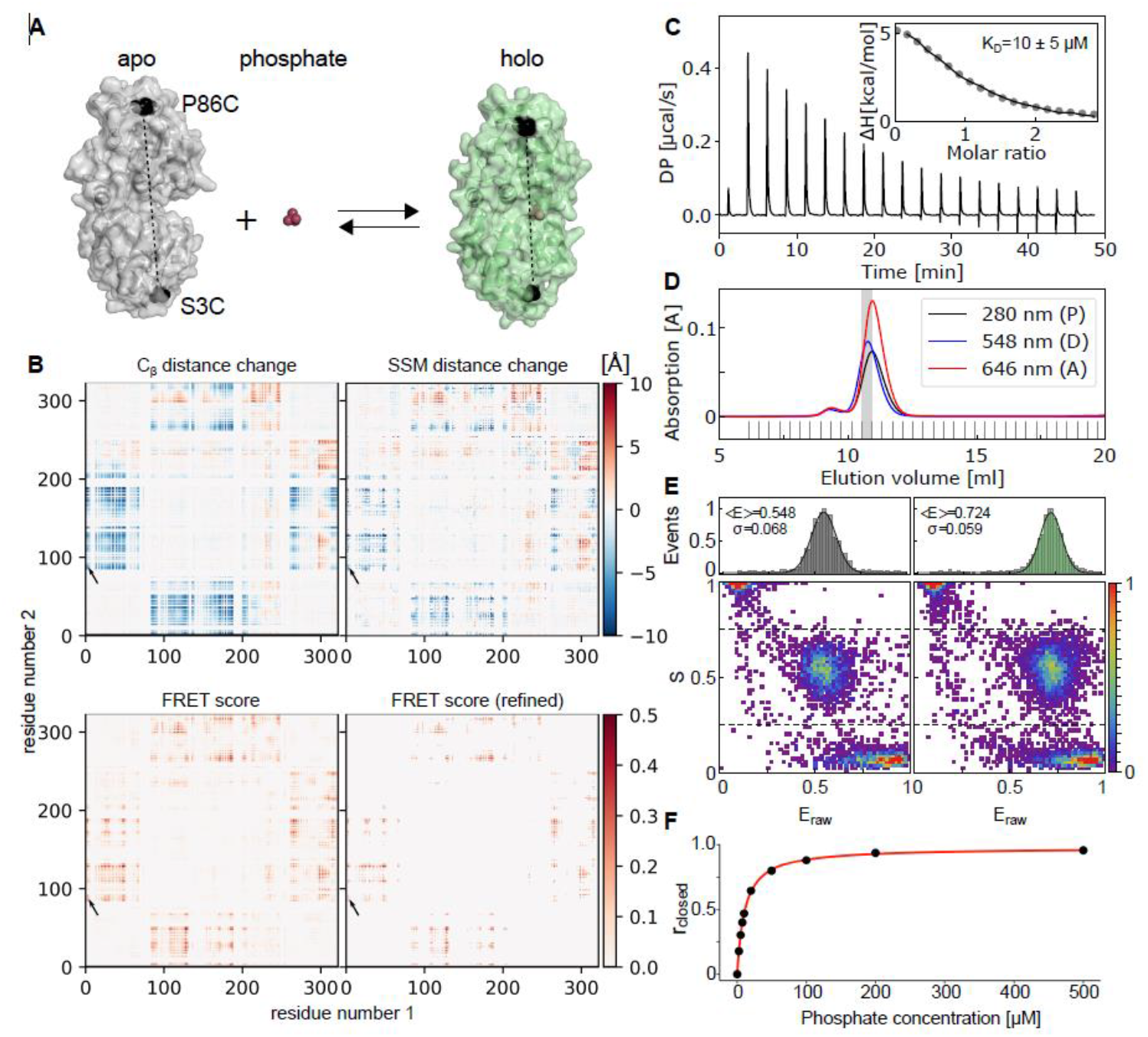
Labelizer-based residue selection for FRET experiments and validation. **A)** Crystal structures of PBP in the apo (grey, PDB-ID: 1OIB) and holo (green, PDB-ID: 1PBP) states with mutations S3C and P86C indicated in black. This variant of PBP also contains an I76G mutation that lowers the affinity for inorganic phosphate by ~200-fold compared to the wild-type protein. **B)** Maps illustrating the distance change and associated FRET score for all pairs of mutants in PBP. The selected mutation S3C and P86C is marked with an arrow. The distance change of the attachment atom (C_ß_, top-left) and the change of the simulated spherical sector model (top-right) show a clear pattern of correlated movements. The calculated FRET score map of all pairs with average label score LS>1 shows only a few spots (~4%) with promising FRET scores FS>0.2 (bottom-left). The selection of the 1000 pairs with highest FRET score and a refinement with FPS software (bottom-right) shows only minor variation compared to the screening map (bottom-left) for the analyzed data points. **C)** Isothermal titration calorimetry (ITC) measurements of PBP (I76G) with average K_d_-values. **D)** Size exclusion chromatography of PBP with LD555-655 showing protein absorbance (280 nm) and fluorophore wavelength at 548 nm and 646 nm. **E)** ES-FRET histograms of PBP (S3C-I76G-P86C) with LD555-655 in the ligand-free apo state (left) and with 480 µM phosphate (right) **F)** Closed fraction (r_closed_) as a function of the substrate concentration for PBP (S3C-I76G-P86C) with Alexa Fluor 555-647 determined from smFRET measurements. The red line is a fitted binding curve; three technical replicates gave an average *K*_D_ of 10.8±0.2 µM.

Subsequently, we labeled all four PBP variants using established procedures^[39,92]^ (see Methods) and studied freely-diffusing molecules with microsecond alternating laser excitation spectroscopy (µsALEX). For labeling, we used the donor-acceptor pair Alexa Fluor 555-Alexa Fluor 647 and the structurally-related combination LD555-LD655 (Figure 6D/E). The success of the labelizer prediction is seen in Figure 6E and Supplementary Figure S9, where high quality smFRET histograms are obtained for all four PBP variants with low FRET in the apo (open conformation, no phosphate) and high FRET in the holo state (closed conformation, 480 µM phosphate). Analyzing the shift of the open to closed conformation and plotting the closed-state fraction as a function of ligand concentrations for PBP (S3C-I76G-P86C) with Alexa555-Alexa647 yields a K_d_ of 16±6 µM (Figure 6F), which is in agreement with results for the unlabeled proteins (Figure 6C). A similar behaviour for the FRET-assay properties and biochemical characteristics are found for all PBP variants (Supplementary Figure S10).

Beside the demonstration of the success of the labelizer procedure, these experiments provide new and so far unavailable information on the ligand binding mechanism of PBP. The lack of a pronounced closed-state population in the absence of ligand (Supplementary Figure S9, apo) and the engulfed nature of the ligand in the closed state support the idea that PBP is likely to use a ligand binding mechanism like other structurally-related SBPs^[58,59,77]^.

To go beyond a qualitative assessment of the labelizer routine, we analyzed a large pool of smFRET experiments of different MalE double-cysteine variants to quantitively benchmark the scores *FS* and *FS*_Δ_. In detail, we analyzed 34 data sets of published^[39,81]^ and unpublished data (Supplementary Figure S11/S12). For these the accurate FRET efficiencies of MalE in both apo- and holo-state are determined including the respective interprobe-distances and their distance change upon maltose binding (Supplementary Data). This data set covers an experimental interprobe distance range from 3-7 nm for three distinct dye pairs with Förster radii of 5.1 nm (Alexa Fluor Alexa-555 Fluor 647), 5.8 nm (ATTO532-ATTO643) and 6.5 nm (Alexa Fluor 546-Alexa Fluor 647)^4^ and *E* values ranging from 0.2-0.9. Consistent with expectations, the calculated *FS* values correlate linearly with the difference of the experimentally determined mean FRET efficiency from 0.5 (Figure 5E). Similarly, we observe a linear correlation between computed *FS*_Δ_ values and the experimentally observed change in FRET efficiency |E_holo_-E_apo_| upon ligand binding (Figure 5E). Whereas pairs with large *FS* and *FS*_Δ_ values are desirable to detect changes upon ligand binding, pairs with high *FS* values of the two individual conformations but *FS*_Δ_ ≈ 0 (MalE 84/352, Supplementary Figure 12) can provide an important experimental control. Such pairs have a distance close to the Förster radius with (almost) no change in FRET efficiency upon conformational change. They can serve as negative controls to ensure that a protein or conformational changes do not influence fluorophores, e.g., via altered photophysics, lifetime and quantum yield changes, or for the characterization of quenchers such as metal ions^[93]^, which can affect FRET efficiencies without conformational change.

Importantly, all analyzed fluorophore-labeled MalE variants used for smFRET had LS values >1 and showed maltose affinities that are wildtype-like with K_d_-values around ~1-2 µM (Supplementary Figure S11). Taken together these analyses provide strong support for the idea that the *LS* is a useful indicator to identify residues that (i) allow fluorophore attachment, (ii) preserve protein function and in combination with FS (iii) enable design of FRET assays.

## Discussion and Conclusions

Here, we present a general strategy to identify optimal residues for protein labeling using a naïve Bayes classifier. Based on a literature screening and bioinformatics analysis of 104 proteins with 396 successfully labeled residues, we identified a set of four parameters, which we combined into a label score to quantitatively rank residues according to their suitability as label site. We show using data from the literature and complementary experiments the predictive power of this labeling score and extend the method to systematically select residue pairs for FRET experiments, which we believe can be extended at later stage to consider the specific properties of the label and also other biophysical assays beyond FRET.

To widely disseminate our methodology, we provide a python package called “labelizer”, which implements the analysis of the pdb-structure, label score calculation, and FRET assay scoring. The labelizer analysis routine can be modified and extended, to accommodate specific research questions and to build upon the work presented here. To make the methodology widely available to non-expert users, all key functionalities are available as a webserver with an intuitive and user-friendly interface https://labelizer.bio.lmu.de/. The webserver supports the label score calculation and its use for FRET experiments with default parameters for the most frequently used fluorophores. For this purpose, pdb-files can be loaded automatically and preprocessed from the pdb-database. We further retrieve conservation scores directly from an independent installation of the ConSurf server^[60,61]^ without the need of uploading any information (except when modified or user-specific pdb files should be used). The webserver visualizes the different scores in an interactive 3D structure viewer and provides a table with filter options for customized restrictions upon residue selection. Furthermore, human-readable result files (csv, json) are available for subsequent analysis. With the developed method, we hope to provide scientists in various research fields (biochemistry, molecular biology, bioimaging, high resolution optical microscopy and single-molecule biophysics) with a tool that enables them to systematically design and justify the residue selection.

A challenging aspect of our analysis is the final selection of residues by the user based on the labelizer output. Since this step is decisive for which residues are used in experiments, the selection goes hand in hand with an assessment and interpretation of the LS/FS value distributions of the analyzed protein. It is difficult to define clear threshold values for residues to be excluded based on LS/FS, yet our findings empirically suggest that residues with LS values < 1 are less likely to be useful in experiments. Since the FRET-score values additionally depend on the underlying LS distribution, it is difficult to give general recommendations. We stress that the user of the algorithm should inspect the specific LS/FS distributions for each protein. For the residues ranked highest, we recommend the user to verify this selection with prior (expert) knowledge on the protein. A key question would be whether the highly-ranked residues, i.e., those favored by the labelizer, are known to negatively impact secondary structure, ligand-binding, biomolecular interactions, or protein folding. Additional information might also come from other biophysical approaches such as CD spectroscopy, FTIR, MD simulations or EPR studies considering any information that can help to assess if key residues, which should not be altered are actually (falsely) suggested by our algorithm.

An interesting future direction for further development of the labelizer is to include more parameters (e.g., also fluorophore-dependent ones) with a potential differentiation of residues based on the selected fluorophores related to the specific charge environment on the protein or proximity to specific amino acids, e.g., tryptophane or histidine. We also plan to combine different parameter scores to improve the predictive ability of the labelizer, which might happen within one category, e. g., via simultaneous use of half-sphere exposure (HSE) and relative surface area (RSA) to combine the amino-acid direction and surface area or between categories, e.g., solvent exposure and cysteine resemblance. Furthermore, normal mode analysis (e.g. NMSim webserver^[94,95]^), mutation specific energy analysis (e.g. SDM^[96,97]^), or tailored MD-simulations^[98]^ could be used to identify FRET-residue pairs for analysis of conformational motion when only one protein structure is available. The concept of FRET scores could be also extended towards other fluorescence assay types related to fluorophore quenching^[99,100]^, protein-induced fluorescence enhancement^[101,102]^, and others^[103,104]^. We also envision applying the labelizer approach in related applications, such as EPR-distance measurements, since the methods share similar requirements in regard to the residue selection^[37–39]^.

Another direction for future improvement and extension of the database and the algorithm, would be to revise the available *PS* values by an extended database, where particularly positions with low or no yield of labeling, could be an important new class of information. Such an improved training data set can be obtained via a feedback loop, where researchers supply information on successfully and unsuccessfully labeled residues via a form planned on our website. Unsuccessful results are of particular interest, since negative results are rarely found in the literature (mainly successful results are published), and we were not able to collect enough negative examples from researchers directly. Therefore, we call on the scientific community to use the labelizer and to provide feedback on the approach and on positive and negative results, where labeling of specific residues was successful or failed, respectively. Finally, once a much larger dataset of labeled and non-labeled residues is available, applications of other machine learning procedures (e.g. support vector machine or neural networks) could significantly enhance the predictions.

## Data and code availability

The webserver with an intuitive user interface and default analysis settings is available under https://labelizer.bio.lmu.de/. The software is available as python package “labelizer” as source code under https://github.com/ChristianGebhardt/labelizer. The databases and additional information can be accessed from https://github.com/ChristianGebhardt/labelizer-supplement or from the online version of the paper.

## Acknowledgements

This work was financed by an ERC Starting Grant (ERC-StG 638536 - SM-IMPORT to T.C.), Deutsche Forschungsgemeinschaft (GRK2062 project C03 to T.C., SFB863 projects A11 and A13 to J.L. and T.C.; Sachbeihilfe CO879/4-1 to T.C.), BMBF (KMU innovative “quantum FRET” to T.C.) LMUexcellent, the Center for integrated protein science Munich (CiPSM) and the Center for Nanoscience (CeNS). We thank all members of the Cordes lab for actively testing the labelizer procedure and webserver, in particular Rebecca Mächtel, Alessandra Narducci, Oliver Brix, Leonor Correia, Shirsha Roy and Chuyu Han. We finally thank our colleagues Gregor Hagelücken, Eitan Lerner, Nicole Robb and Giorgos Gouridis for discussions and support of the project.

## Author contributions

C.G. and T.C. conceived and designed the study. C.G performed research, data analysis and software implementation. J.L. provided new analytical tools. K.S. and T.C. analyzed data. C.G., P.B., R.S. and K.S. implemented the webserver. A.K.S., N.W. and G.G.M.M. performed research. D.A.G. prepared PBP variants, performed research and analyzed data. J.L. and T.C. supervised the study. C.G., J.L. and T.C. discussed and interpreted the results and wrote the manuscript in consultation with all authors.

## Online Methods

### Database generation

To identify parameters with predictive power for the possibility to label residues in proteins, we created a dataset based on a non-automated screening of more than 1000 publications published or preprinted which were available on or before December 2020 with a focus on the field of single-molecule microscopy and single-molecule FRET. The papers were screened to identify proteins and residues that were labeled successfully with a fluorophore and that satisfied the following criteria: (i) the proteins had a structure available in the PDB-database (with PDB identification code); (ii) the protein was labeled via site-specific mutagenesis and introduction of cysteines or UAAs; (iii) the protein was successfully labeled synthetic organic fluorophores (or spin labels) and used preferentially single-molecule assays. In order to increase the number of database entries, we complemented our search whenever some information was missing. Typical cases were missing PDB identification codes or residue numbers. In this case, the required information was obtained from other referenced papers (often) of the same research group.

For each successfully labeled protein variant, which fulfilled the aforementioned criteria, the following information was collected^5^:

- Protein (PDB identification code)
- Soluble or membrane protein
- Stoichiometry (monomers, dimer, complexes)
- Homology model (true/false)
- Labeled residue (chain and residue number)
- Mutation (cysteine or UAA)
- Assay type (smFRET, imaging, bulk-FRET, other)
- Name of labeled fluorophores
- Research group
- Publication reference

The final database with information on those positions in proteins that were successfully labeled had 396 successfully labeled residues in 112 different chains in 104 different protein structures (Supplementary Data). As comparison, we used a representative set of proteins (PDBselect, November 2017)^[65,66]^ as a random reference database to check how representative the analyzed pdb structures are. Therefore, we randomly selected 300 chains (out of 4184 chains) from the PDBselect database and performed the identical analysis with those pdb files. This important comparison shows that the selection of labeled proteins and residues is representative of the pdf content, indicated by only minor deviations between both P(s) distributions, mostly within statistical errors (see Supplementary Figure S2).

#### Parameter frequency calculation

For every extracted parameter, the relative frequency defines a parameter score

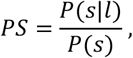

where *P*(*s*) is the probability distribution of the score *s* (calculated from the 112 chains of the database) and *P*(*s*|*l*) is the probability distribution of the score given that the residue was labeled (calculated from the 396 successfully labeled residues).

The error bars *σ*_s*l*_ and *σ*_s_ for *P*(*s*|*l*) and *P*(*s*), respectively, were determined from Poissonian counting statistics as 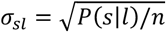 and 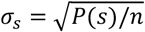 with *n* being the total number of evaluated residues. The error bar *σ*_*PS*_ for *PS* follows from standard error propagation rules:

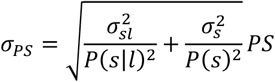

#### Parameter information analysis

To evaluate the amount of information a single parameter score inheres, we used three measures to estimate the deviation from an equal distribution, which corresponds to the case of zero information.

We used standard Pearson correlation for a pair of numeric parameters

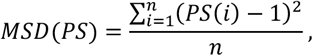

with n the number of bins/categories.

We used standard Pearson correlation for a pair of numeric parameters

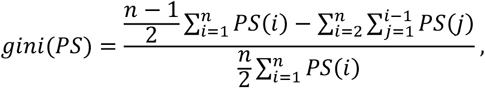

with n the number of bins/categories.

We used an adapted Shannon entropy accounting for the number of bins/categories as

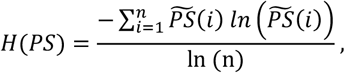

with a normalized parameter score 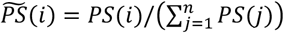 and n the number of bins/categories.

#### Parameter cross-correlation

To evaluate the mutual statistical dependence of all calculated parameters, we use three different types of correlation coefficients, depending on the datatypes of the parameters:

We used standard Pearson correlation for a pair of numeric parameters

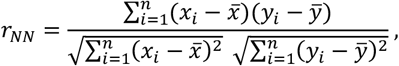

with *n* different residues with parameter scores *x*_*i*_, *y*_*i*_ and corresponding mean values 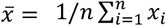 (and 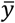 accordingly)^[105]^.

We used the interclass correlation for a pair of a categorical parameter and a numeric parameter^[106]^. The *n* data points are grouped in k categories *c*_*i*_ *with i* ∈ {1,2, …, *k*} of length *n*_*i*_.

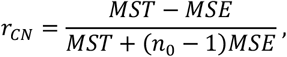

with

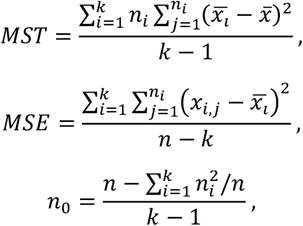

where 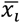 is the mean of category *i*, 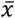 the mean of all data, *x*_*i,j*_ the j^th^ numeric value in category *c*_*i*_, and (*n*_0_ − 1) the averaged interclass degree of freedom^[106]^.

We used the Cramer’s V for a pair of a categorical parameters^[107]^. The data are grouped in the two categories *c*_*i*_ *w*i*th i* ∈ {1,2, …, *k*} and *d*_*j*_ *with j* ∈ {1,2, …, *l*}.

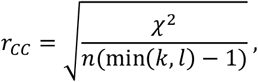

with

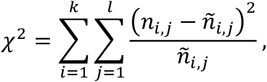

where 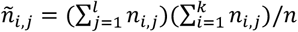, *n* total number of residues and *n*_*i,j*_ number of residues of class *c*_*i*_ and *d*_*j*_. The cross-correlation was calculated for every combination of the 28 extracted parameters to identify dependencies as shown in Figure 2.

#### Parameter Selection Criteria

The selection of a suitable parameter set is based on two criteria. First, a joined correlation for any combination of parameters is calculated as

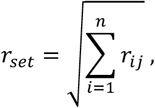

with *r*_*ij*_ the correlation of parameter *i*with *j* and *n* the number of selected parameters (in our case 4). Secondly, we used three measures to characterize our parameter sets:

We calculate the t value of the calculated label scores as

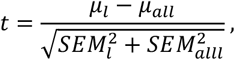

with the mean values *μ*_l_, *μ*_*all*_ and standard error of mean *SEM*_l_, *SEM*_*all*_ of the labeled/all residues, respectively.

The dynamic range was calculated as the standard deviation of the logarithmic values *σ*(log(*LS*_*all*_)).

The suppression/enhancement of the labeling score of labeled residues for small/large values was calculated from the slope of a linear least square fit to the logarithm of the label score *LS* and the label score distribution of labeled residues and all residues. The data are binned into logarithmic bins with bin intervals [1.5^*i*^, 1.5^*i*+1^] *for i* ∈ {−12, …, 11} and fitted to the function

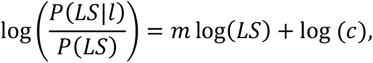

where LS is the label score and *P*(*LS*)/ *P*(*LS*|*l*) the probability distributions of the label score of all and the labeled residues. The slope *m* is used as analysis parameter form the fitted values *m, c*.

### Protein production and labeling

In the current study we used single cysteine variants of MalE (Figure 4) that were obtained and fluorophore-labeled as described previously^[59,108]^. PBP double cysteine variants were produced for this study. The coding sequence for the *E. coli* K12 *phoS* gene (Genbank coding sequence NC_000913.3, 3910485 - 3911525 complement, protein accession number NP_418184.1), with amino-acid changes (A17C and A197C) corresponding to the rho-PBP fluorescent biosensor variant^[109]^ was synthesized (Invitrogen GeneArt Gene Synthesis, Thermo Fisher) without its N-terminal signal sequence (25 amino acid N-terminal deletion). This construct utilized flanking NdeI/XhoI sites, and was subcloned into the pET20b expression vector. The resulting construct encoded a C-terminal His-tag fusion. The S3C-P86C-PBP mutant, with the additional I76G mutation that reduces the wild-type affinity (*K*_*d*_ 0.07 µM) of the protein for inorganic phosphate by ~200-fold^[91]^ was created using a protocol based on the Stratagene Quikchange protocol^[110]^. As a control, a variant was also created with only the I76G mutation.

*E. coli* BL21 (DE3) *pLysS* cells transformed with the S3C-P86C-PBP mutant expression plasmid (or the plasmid for the control variant) were used to inoculate Terrific Broth (TB; Carl Roth, Karlsruhe, Germany) supplemented with 100 µg/ml carbenicillin (Carl Roth) and 0.2% glucose to an optical density at 600 nm (OD_600_) of 0.1 AU at 37°C with shaking at 200 rpm. At an OD_600_ of ~0.3 AU, isopropyl b-D-1-thiogalactopyranoside (IPTG, Carl Roth) was added to a final concentration of 0.5 mM, followed by ~24 h incubation. Cells were harvested by centrifugation (5000*g*, 20 min, 4°C) at a final culture OD_600_ of 3-4 AU, resuspended in 35 ml 20 mM HEPES pH 7.5, 300 mM NaCl, 10% glycerol containing protease inhibitor (cOmplete, EDTA-free Protease Inhibitor Tablets, Sigma; 1 tablet/50 ml solution), and frozen and stored at −80°C.

The resulting cell suspension was thawed, supplemented with 5 mM β-mercaptoethanol (β-ME) and 10 mM imidazole (Carl Roth), and then sonicated (Branson Digital Sonifier 450, Danbury, CT, USA) on ice for 10 min (Amplitude, 25%; 0.5 sec on and 0.5 sec off). Insoluble fractions containing cell debris were separated by centrifugation (165,000*g* for 1 h at 4°C). The soluble fraction was incubated with 1.5 ml of Ni Sepharose 6 Fast Flow resin (GE Healthcare) for 1 h at 4°C. The resin with bound protein was then washed with 80 ml of buffer containing 25 mM imidazole. Bound protein was eluted in 10 ml buffer with 500 mM imidazole. The elution fraction was concentrated to < 0.5 ml using a Viva Spin 20 concentrator with a 10 kDa MWCO (Th. Geyer, Renningen, Germany), and subjected to further purification by size-exclusion chromatography (SEC; using ÄKTA pure system, and Superdex 75 Increase 10/300 GL column (GE Healthcare)) in 20 mM Tris-HCl pH 8.0, 100 mM NaCl, 10 mM imidazole. The final purified proteins were >95% pure as assessed by sodium dodecyl sulfate polyacrylamide gel electrophoresis (SDS-PAGE).

His-tagged MalE, and S3C-P86C-PBP proteins were labeled as described previously^[59,108]^. The proteins were incubated with 1 mM DTT to reduce cysteine residues. Following dilution to lower the DTT concentration to < 0.05 mM (so as not to interfere with binding of protein to the metal-affinity resin), the proteins were immobilized on 200 µl of Ni Sepharose resin. The resin was then washed with 12 ml of 50 mM Tris-HCl pH 7.4-8.0, 50 mM KCl, 5% glycerol for MalE and SBD2 (Buffer A), and 20 mM Tris-HCl pH 8.0, 100 mM NaCl, 10 mM imidazole for PBP. 28 nmoles of PBP were then incubated overnight with 50 nmol of each fluorophore dissolved in 2 ml of the appropriate buffer. Unreacted fluorophore for MalE and SBD2 was removed by washing the resin with 12 ml of Buffer A followed by 12 ml of Buffer A containing 50% glycerol. For PBP, a single 12 ml wash was performed. Bound MalE and SBD2 were eluted with 0.5 ml of Buffer A containing 500 mM imidazole, whereas PBP was eluted with 1 ml of buffer with 500 mM imidazole. The labeled proteins were further purified by size-exclusion chromatography (using ÄKTA pure system, and Superdex 75 Increase 10/300 GL column (GE Healthcare)). Absorbance of protein (280 nm) and fluorophore (488 nm, 532 nm and 640 nm) was used for determination of molar concentrations in samples and labeling efficiency, i.e., [Fluorophore]/[protein]*100.

### Affinity measurements: Isothermal titration calorimetry and MST

Binding affinities of I76G-PBP and unlabeled S3C-P86C-PBP for inorganic phosphate were determined with a MicroCal PEAQ-ITC microcalorimeter (Malvern Panalytical) at 25°C. Protein from a diluted solution was concentrated to ~ 30 µM using a Viva Spin 6 concentrator with a 10 kDa MWCO. The filtrate was used to prepare the phosphate solution at 450 µM. The reaction cell was filled with the protein solution and titrated in 19 steps of 2 µl each of phosphate solution in 160-s intervals. A baseline control was obtained from measurements made with protein-free filtrate in the reaction cell, and this baseline was subtracted from the experimental thermograms. Data were fitted to a single binding site model using the Setup MicroCal PEAQ-ITC Analysis Software provided by the manufacturer.

### smFRET spectroscopy and data analysis

smFRET experiments of PBP were carried out on a home-built ALEX setup as described previously^[58,92]^: PBP was studied by diluting the labeled protein to concentrations of ≈80 pM in a 100 µl drop of buffer (20 mM Tris-HCl pH 8.0, 100 mM NaCl, 10 mM imidazole) on a coverslip supplemented with the ligand phosphate as described in the text and figures. Before each experiment, the coverslip was passivated for 3 minutes with a 1 mg/ml BSA solution in buffer. The measurements were performed without photostabilizer. The fluorescent donor molecules were excited by a diode laser at 532 nm operated at 60 µW at the sample in alternation mode (50 µs alternating excitation and a 100 µs alternation period). The fluorescent acceptor molecules were excited by a diode laser at 640 nm operated at 25 µW at the sample. Data analysis was performed using a home written software package as described in reference ^[58]^. Single-molecule events were identified using an all-photon-burst-search algorithm with a threshold of 15, a time window of 500 µs and a minimum total photon number of 150^[111]^. E-histograms of double-labeled FRET species with LD555 and LD655 were extracted by selecting 0.25<S<0.75. E-histograms of the open state without ligand (apo) and closed state with saturation of the ligand (holo) were fitted with a Gaussian distribution 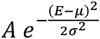.

## Supporting Information

### A) Supplementary Note 1: Database parameter evaluation

#### Data preprocessing

the 104 PDB files of the database and the comparison PDB files were downloaded from the protein databank and preprocessed to unify the data structures. Therefore, all hetero atom entries (HETATM), anisotropy entries (ANISOU), and connection entries (CONNECT, as well as all the meta-information (REMARK) were removed from the pdb files^1,2^. Chains of polymeric protein assemblies in crystals were deleted if these were bare crystallization artifacts and do not occur in natural environments. The conservation score was calculated for all 112 chains containing the labeled residues (and the reference database) with the default settings (see Supplementary Table S1)^3,4^. Failed conservation score calculations (e.g. if too few homologue structures are available) were ignored for further analysis.

### PDB data processing

The pdb files are parsed and processed with the “Bio.PDB” module^5^ of the “biopython” package^6^.

### Parameter extraction

28 parameters were calculated or extracted from third party software and assigned to the four categories (i) solvent exposure, (ii) residue conservation, (iii) cysteine resemblance, and (iv) secondary structure (see Supplementary Table S1-4).

Overall, 43357 and 29898 residues from the database and reference dataset are considered in the calculations, respectively. Failed parameter calculations were ignored for further analysis. Therefore, the number of calculated values varies for the 28 parameters (failure rate <10% for all parameters in the database and reference database; see Supplementary Table S5 for settings of the ConSurf-server and Supplementary Table S6 for exact numbers).

**Supplementary Table S1.**
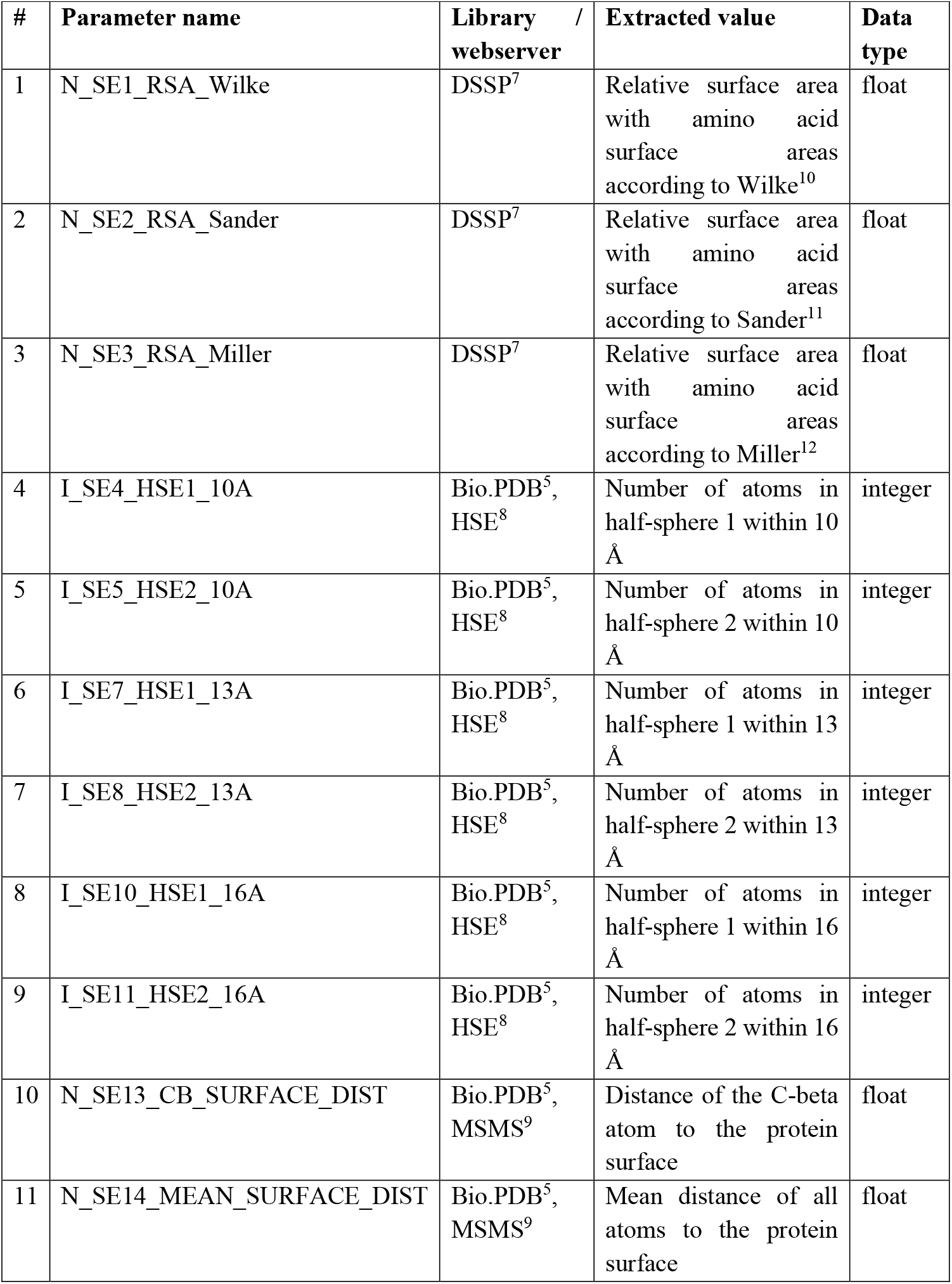
Solvent exposure related values were extracted using the third party algorithms (i) “Define Secondary Structure of Proteins” (DSSP) to calculate relative surface accessibility^7^, (ii) “Half-Sphere-Exposure” (HSE) to calculate the number of C-alpha atoms in the half-spheres defined by the C-alpha – C-beta vector^8^, and (iii) “Michel Sanner’s Molecular Surface” (MSMS) to calculate the protein surface and the residue depth of the atoms^9^.

**Supplementary Table S2.** Parameters related to residue conservation are extracted from the grades-file of the consurf server^3,4^. Settings for the ConSurf analysis are listed in Table S5.

| # | Parameter name | Library /<br>webserver | Extracted value | Data type |
| --- | --- | --- | --- | --- |
| 12 | I_CS1_Color | ConSurf<sup>3,4</sup> | Color representation of conservation score (binned conservation score with upper and lower boundary) | integer |
| 13 | N_CS2_Score | ConSurf<sup>3,4</sup> | Conservation score | float |
| 14 | N_CS3_Lower_Score | ConSurf<sup>3,4</sup> | Lower value of confidence interval of conservation score | float |
| 15 | N_CS4_Upper_Score | ConSurf<sup>3,4</sup> | Upper value of confidence interval of conservation score | float |
| 16 | I_CS5_Variety_Length | ConSurf<sup>3,4</sup> | Number of different amino acids in homologues | integer |
| 17 | C_CS6_Cys_In_Variety | ConSurf<sup>3,4</sup> | Boolean: true if cysteine is in amino acid list of homologues; false else | categorical |

**Supplementary Table S3.** Secondary structure related values were extracted using the third party algorithms “Define Secondary Structure of Proteins” (DSSP) to calculate the secondary structure of the residue of interest and its adjacent residues as well as the backbone torsion angles^7^.

| # | Parameter name | Library /<br>webserver | Extracted value | Data type |
| --- | --- | --- | --- | --- |
| 18 | C_SS1_SS | DSSP<sup>7</sup> | Secondary structure | categorical |
| 19 | N_SS2_Phi | DSSP<sup>7</sup> | Backbone torsion angle (n-1)-n | float |
| 20 | N_SS3_Psi | DSSP<sup>7</sup> | Backbone torsion angle n-(n+1) | float |
| 21 | C_SS4_SS-1 | DSSP<sup>7</sup> | Secondary structure of predecessor residue | categorical |
| 22 | C_SS4_SS-2 | DSSP<sup>7</sup> | Secondary structure two positions before | categorical |
| 23 | C_SS4_SS+1 | DSSP<sup>7</sup> | Secondary structure of successor residue | categorical |
| 24 | C_SS4_SS+2 | DSSP<sup>7</sup> | Secondary structure two positions after | categorical |

**Supplementary Table S4.**
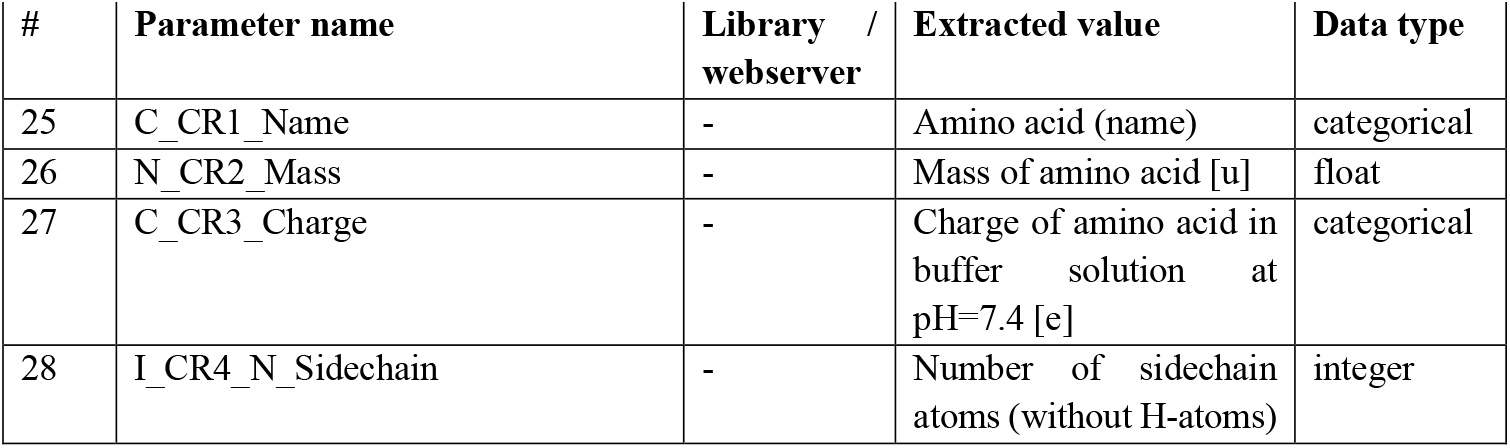
Cysteine resemblance related values were taken from the amino acids structures to either compare the individual amino acids or group the amino acids by charge and size/mass.

**Supplementary Table S5.** ConSurf-server settings. Overview of all user parameters set for the conservations score calculation on https://consurf.tau.ac.il/ (accessed January 24^th^, 2021).

| Parameter | Value |
| --- | --- |
| DNA_AA | AA |
| NMR | no |
| PDB_yes_no | yes |
| MSA_yes_no | no |
| Homolog_search_algorithm | HMMER |
| ITERATIONS | 1 |
| E_VALUE | 0.0001 |
| proteins_DB | UNIREF90 |
| user_select_seq | no |
| MAX_NUM_HOMOL | 150 |
| best_uniform_sequences | uniform |
| MAX_REDUNDANCY | 95 |
| MIN_IDENTITY | 35 |
| MSAprogram | MAFFT |
| ALGORITHM | Bayes |
| SUB_MATRIX | BEST |

**Supplementary Table S6.** Parameter overview and statistics. The table summarizes the number of analyzed residues (complete database, labeled residues) and the statistical evaluation of the parameter scores with respect to mean-square deviation (MSD), gini coefficient, and adapted Shannon entropy (see methods for details).

| # | Name | Number of all analyzed residues | Number of labeled residues | MSD | Gini coeff | Shannon entropy |
| --- | --- | --- | --- | --- | --- | --- |
| 1 | N_SE1_RSA_Wilke | 40056 | 385 | 0.951 | 0.303 | 0.931 |
| 2 | N_SE2_RSA_Sander | 40056 | 385 | 0.866 | 0.323 | 0.923 |
| 3 | N_SE3_RSA_Miller | 40056 | 385 | 0.948 | 0.330 | 0.922 |
| 4 | I_SE4_HSE1_10A | 40056 | 385 | 0.674 | 0.663 | 0.687 |
| 5 | I_SE5_HSE2_10A | 40056 | 385 | 0.259 | 0.421 | 0.851 |
| 6 | I_SE6_HSE1_13A | 40056 | 385 | 0.922 | 0.625 | 0.712 |
| 7 | I_SE7_HSE2_13A | 40056 | 385 | 0.266 | 0.385 | 0.861 |
| 8 | I_SE8_HSE1_16A | 40056 | 385 | 0.972 | 0.596 | 0.744 |
| 9 | I_SE9_HSE2_16A | 40056 | 385 | 0.206 | 0.347 | 0.895 |
| 10 | N_SE10_CB_SURFACE_DIST | 40056 | 385 | 0.210 | 0.585 | 0.721 |
| 11 | N_SE11_MEAN_SURFACE_DIST | 40056 | 385 | 0.883 | 0.578 | 0.744 |
| 12 | I_CS1_Color | 39409 | 376 | 0.322 | 0.271 | 0.940 |
| 13 | N_CS2_Score | 39409 | 376 | 0.856 | 0.339 | 0.914 |
| 14 | N_CS3_Lower_Score | 39409 | 376 | 0.869 | 0.469 | 0.819 |
| 15 | N_CS4_Upper_Score | 39409 | 376 | 0.530 | 0.343 | 0.900 |
| 16 | I_CS5_Variety_Length | 39409 | 376 | 0.677 | 0.354 | 0.911 |
| 17 | C_CS6_Cys_In_Variety | 39409 | 376 | 0.006 | 0.038 | 0.996 |
| 18 | C_SS1_SS | 41502 | 390 | 0.218 | 0.307 | 0.894 |
| 19 | N_SS2_Phi | 41502 | 390 | 0.448 | 0.370 | 0.869 |
| 20 | N_SS3_Psi | 41502 | 390 | 0.231 | 0.339 | 0.883 |
| 21 | C_SS4_SS-1 | 41227 | 385 | 0.258 | 0.383 | 0.861 |
| 22 | C_SS5_SS-2 | 41018 | 382 | 0.175 | 0.250 | 0.919 |
| 23 | C_SS6_SS+1 | 41227 | 388 | 0.218 | 0.316 | 0.900 |
| 24 | C_SS7_SS+2 | 41021 | 385 | 0.207 | 0.259 | 0.917 |
| 25 | C_CR1_Name | 43357 | 396 | 0.753 | 0.401 | 0.909 |
| 26 | N_CR2_Mass | 43357 | 396 | 0.361 | 0.367 | 0.905 |
| 27 | C_CR3_Charge | 43357 | 396 | 0.156 | 0.162 | 0.956 |
| 28 | I_CR4_N_Sidechain | 43357 | 396 | 0.413 | 0.368 | 0.902 |

### B) Supplementary Note 2: Förster resonance energy transfer

#### Förster radius calculation

Spectral information, quantum yield, and extinction coefficients are taken from the database https://www.fpbase.org/spectra/ and provided with the labelizer-package for the most commonly used fluorophores.

The Förster radius *R*_*0*_ is given by

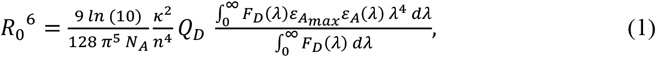

whereby *Q*_*D*_ is the donor quantum yield, *F*_*D*_ the normalized donor emission spectrum, *ε*_A_ the normalized acceptor emission spectrum, and 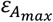 the acceptor extinction coefficient.

The following values are set fix to theoretical values:

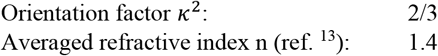

#### Distance screening

Center of mass of a sphere with radius R cut with a cone of angle *α* in the z-dimension (spherical sector):

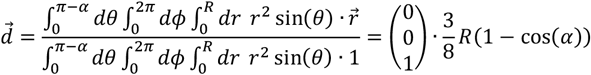

We assume the origin to be at the attachment site (C-ß atom) of the fluorophore and model the accessible volume of the dye with a cut sphere of angle *α* and the non-accessible space with *π* − *α*. We approximate the center of mass of the non-accessible volume 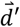 with the center of mass of all *N* atom positions 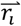 within the protein closer than *R* to the attachment point (see Figure 5):

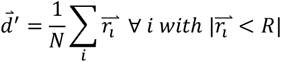

The direct relation between the center of mass of atoms in the protein 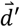 and the center of mass of the fluorophores accessible-volume 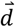 is given by:

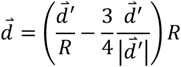

We add an empirically determined correction factor based on simulations with 35 different fluorophore parameters on 100 residue pairs in 10 different protein structures to account for the finite size of atoms and fluorophores, which leads to a gap between protein atoms and accessible volume (see Supplementary Figure S7B/D).

The offset for the protein surface is added as a small addition to the fluorophore linker

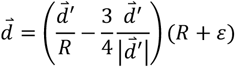

and reads as

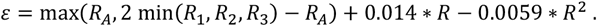

This correction can reproduce the simulated mean positions 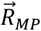 with a root mean square deviation of ±2.7 Å (see Supplementary Figure S7F) and mean position distances 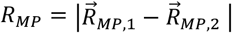 with a root mean square deviation of ±2.1 Å (see Figure 8C).

We approximated the relation between the mean positions of the accessible volumes to the measured FRET averaged distances as

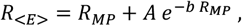

whereby the second term accounts for FRET-efficiency weighted averaging effects at small distances (see Supplementary Figure 8B). The values *A* and *b* are determined as *A* = 20.6 Å and *b* = 0.037 1/Å from a fit to the 35000 simulated distances within the ten selected protein structures, which is similar to the reported relations in ref. ^14^ and ^15^ for DNA. With this relationship, the simulated distance with FPS is reproduced up to a deviation of ±3.4 Å (±3.1 Å for distances between 40 and 75 Å).

Based on the corrected FRET values, the (screening) FRET-efficiency of a residue pair {i,j} is calculated as

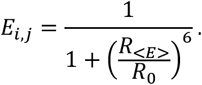

#### Distance refinement

A refinement is calculated based on the N highest FRET scores (with default N=300) using the available FPS simulation software^16^ with standard parameter settings (see Supplementary Table S8).

### C) Supplementary Figures

**Supplementary Figure S1.**
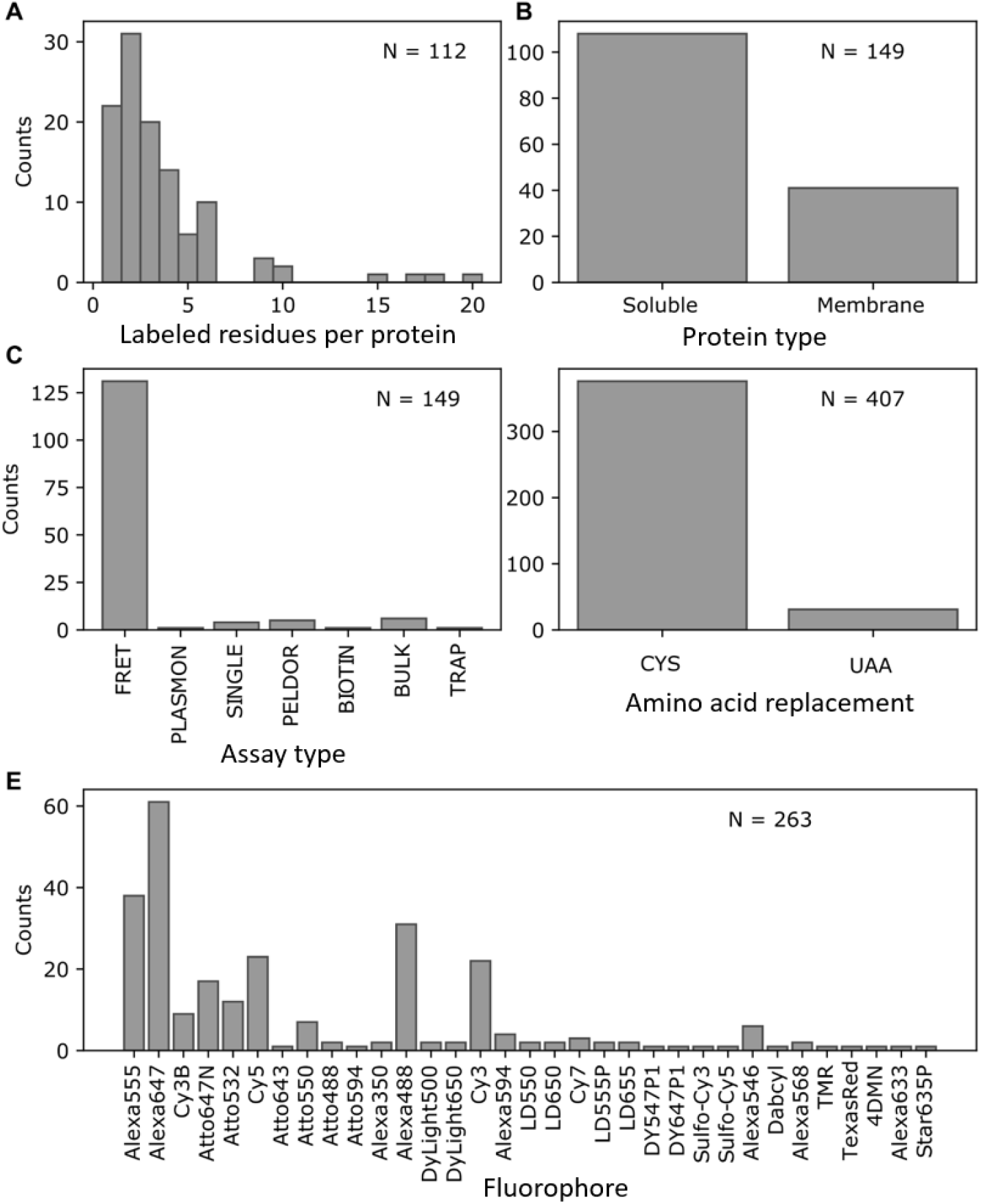
Labeling database statistics. **A)** Number of labeled residues per chain in the database with N=112 different protein chains. **B)** Comparison of published protein systems with soluble and membrane proteins (N=149 published protein systems, multiple occurrence possible). **C)** Statistics of the different assay types used for the labeling database (N=149 published protein systems, multiple occurrence possible). Around 90% of the assays are single-molecule FRET assays (FRET), the others are bulk FRET (BULK) or single fluorophore labeled (SINGLE) assays, spin labels (PELDOR), gold labels (PLASMON), biotin labels (BIOTIN) and linker labels for optical traps (TRAP). **D)** Statistics on the labeling residues (cysteine or unnatural amino acid, N=407 residues). **E)** Statistics on the fluorophores used in the publications (N=263 occurrences in the publications).

**Supplementary Figure S2.**
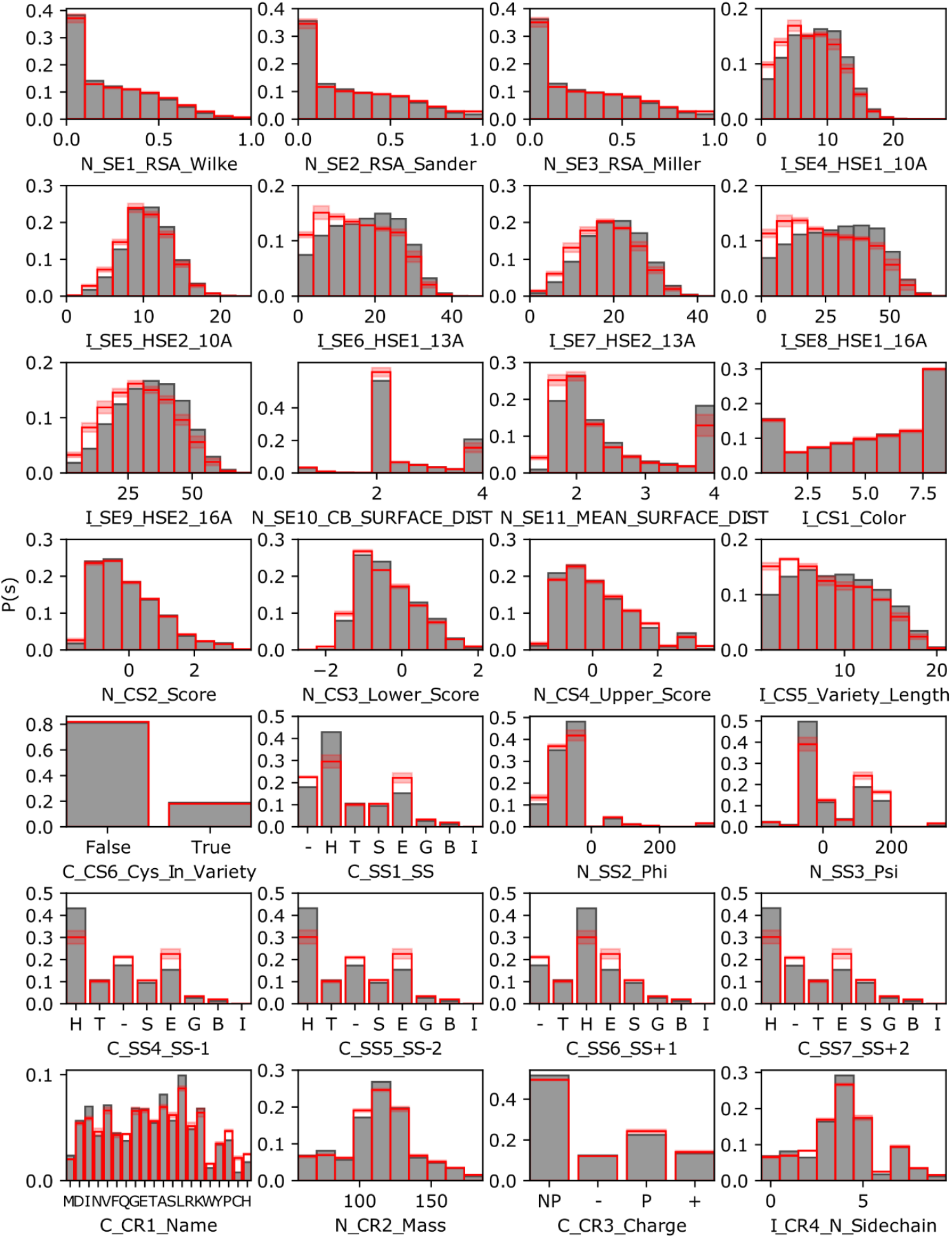
Parameter distribution comparison. Probability distributions *P*(*s*) of all 28 scores in the labeling database (gray). The values are compared to a randomly selected representative reference dataset (red line: mean values, pale area: standard deviation of triplicates) based on the pdbselect dataset^17,18^ (see methods for details). The x-axis label specifiers the parameter: first part for the type of data (I: integer, N: numeric, C: categorical), second part for the parameter group (SE: solvent exposure, CS: conservation score, SS: secondary structure, CR: cysteine resemblance), and the rest for a reasonable name (see methods for details).

**Supplementary Figure S3.**
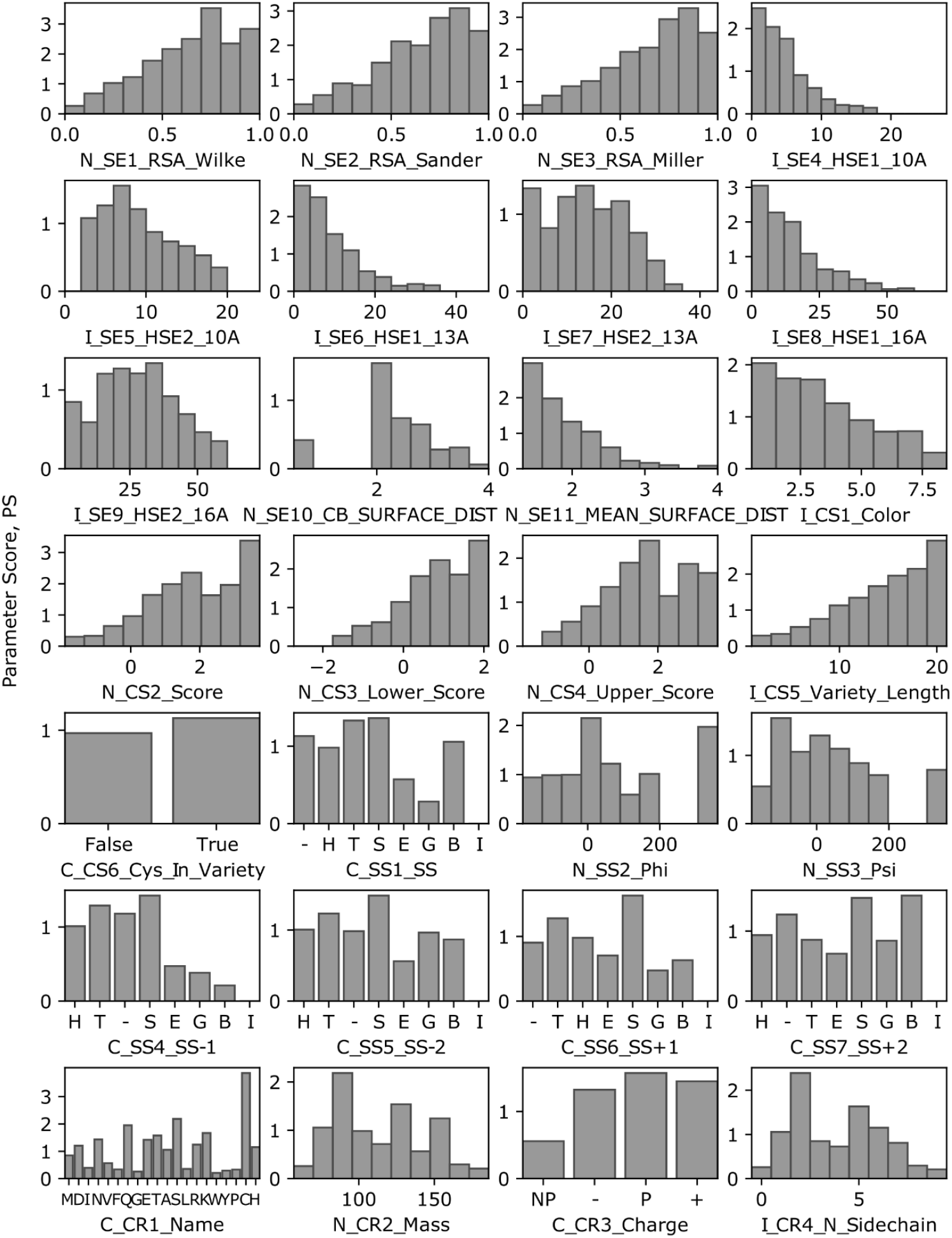
Conditional frequencies of parameters. Conditional frequency distributions *P*(*s*|*l*)/P(*s*) defining the parameter scores of all 28 parameters in the labeling database. The x-axis label specifiers the parameter: first part for the type of data (I: integer, N: numeric, C: categorical), second part for the parameter group (SE: solvent exposure, CS: conservation score, SS: secondary structure, CR: cysteine resemblance), and the rest for a reasonable name (see methods for details).

**Supplementary Figure S4.**
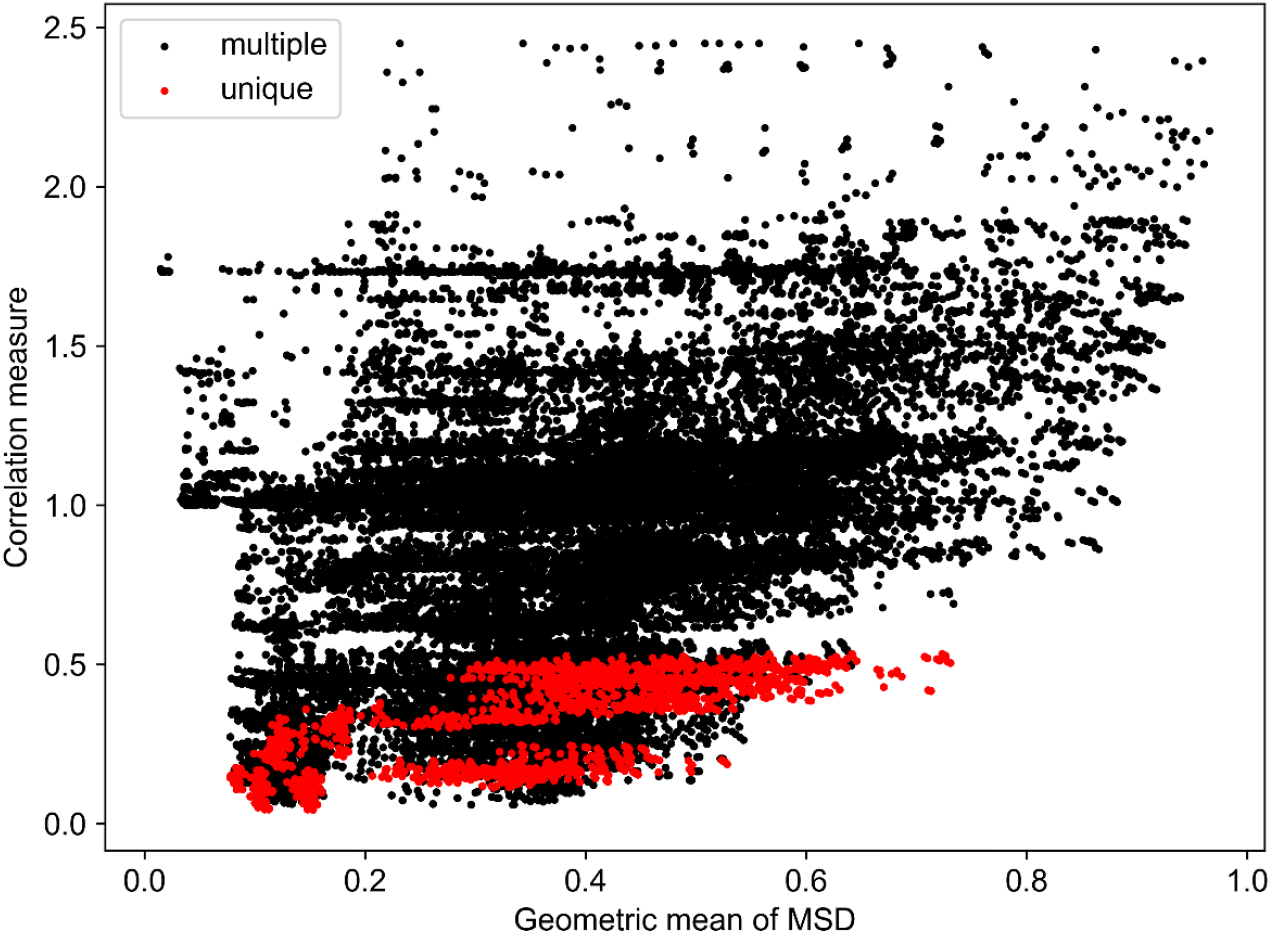
Correlation measure and averaged mean square deviation (MSD). The plot shows the geometric mean of the MSD value of the selected parameters (mean square deviation from equal contribution, see methods) versus the correlation measure (2-norm of all correlations). All parameters are combined with each other to sets of 4, whereby points with multiple (2 or more) parameters from the same group (e.g. solvent exposure) are marked black. Combinations with one parameter from each group are marked red.

**Supplementary Figure S5.**
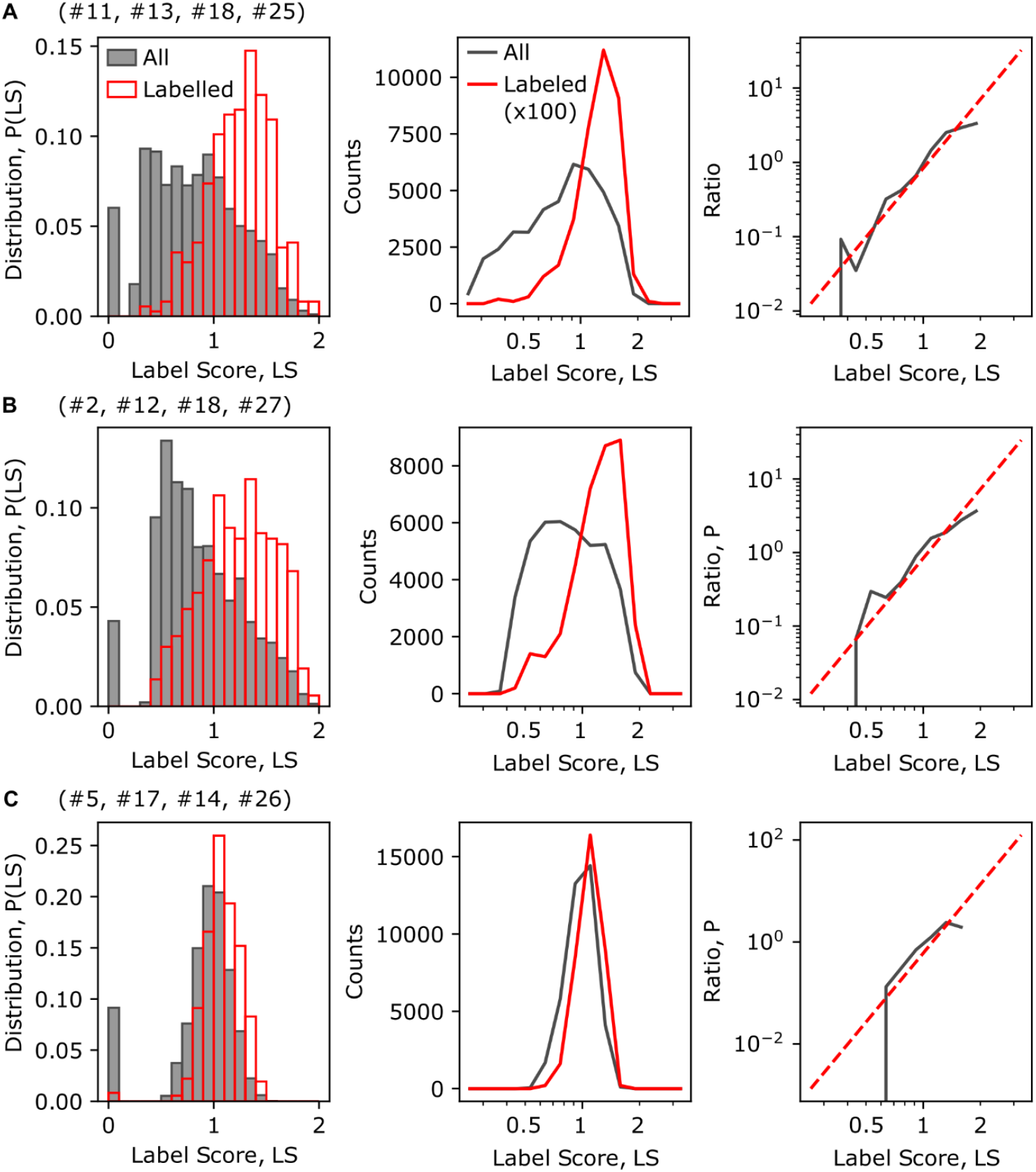
Label score evaluation for different parameter sets. **A** Label score probability distribution of all residues (gray) and labeled residues (red) in our database (left) and the a histogram with logarithmic scale of the label scores (middle) for the selected quadruple (#11: mean surface distance, #13: ConSurf score, #18: secondary structure, #25: amino acid identity) (default settings). The ratio of the probability distribution of labeled and all residues (gray) is fitted with a linear dependency (red, dashed) in the log-log-plot (right). **B** Same evaluation as in A for another suitable parameter selection of the quadruple (#2: relative surface area Sander, #12: ConSurf color, #18: secondary structure, #27: amino acid charge). **C** Same evaluation as in A for a parameter set with poor prediction power (#5: second half of 10 Å half-sphere exposure, #17: cysteine in homologue structures, #14: secondary structure two positions after, #26: amino acid mass).

**Supplementary Figure S6:**
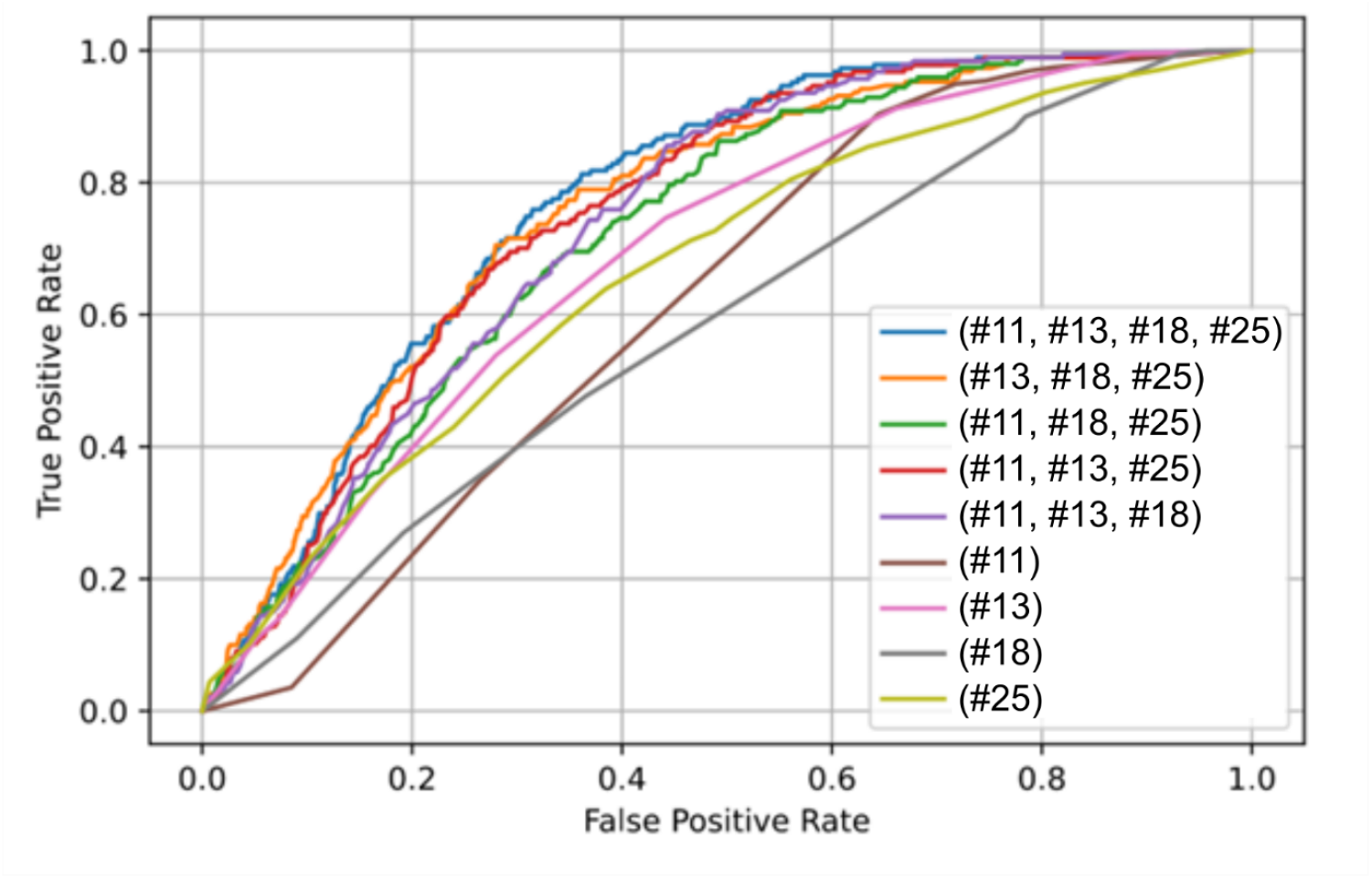
ROC curves for reduced parameter sets. To study the importance of individual parameters in the final prediction, we compared the receiver operating characteristic (ROC curve) for the baseline, when removing one of four parameters and for each parameters on its own. We considered labeled sites positives and unlabeled site negatives, which most likely overestimates the false-positive rate.

**Supplementary Figure S7.**
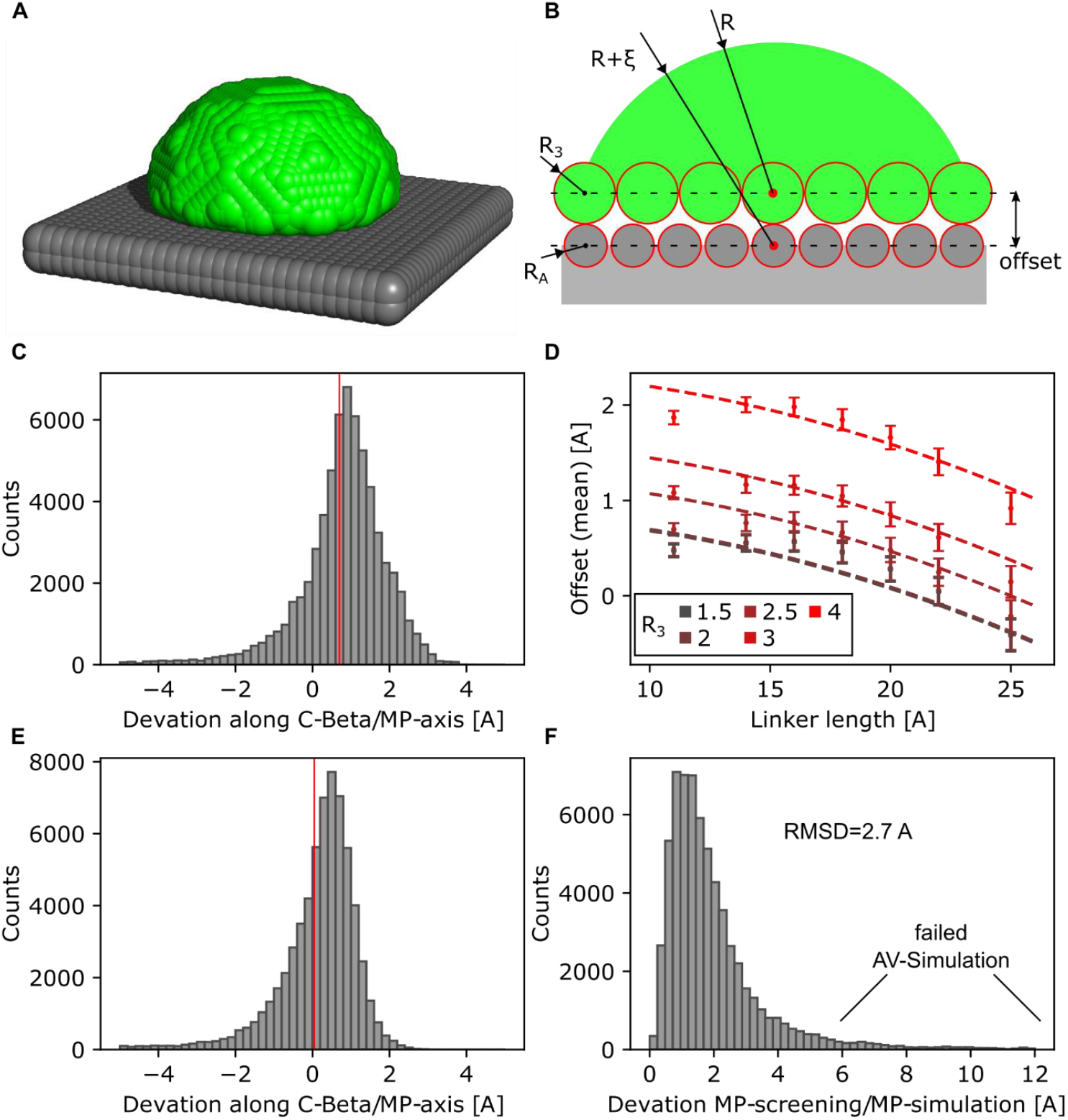
Correction parameter for mean position of accessible volume. **(A)** Simplified accessible volume simulation (green) in an idealized system of a planar array of atoms (gray). **(B)** Motivation for the correction factor is illustrated with the offset between the atom coordinates (lower dashed line) and the accessible volume coordinates (upper dashed line). The correction factor *ε* corrects for this gap between accessible surface (green) and inaccessible surface (gray) under the consideration of the linker length (R, corresponds to the AV radius), the atom radius *R*_*A*_ and the smallest fluorophore radius *R*_3_ of the ellipsoidal approximation^14,16,19^. **(C)** Deviation between simulated mean position of accessible volume (FPS software) and estimated mean position (SSM approach) with indicated mean value (red line). **(D)** Mean offset from (C) for different linker lengths R and fluorophore radii R_3_ is shown with error bars (standard error of the mean from simulations). The estimation of the offset in (C) is fitted globally with the correction factor *ε* = max(*R*_*A*_, 2 min(*R*_1_, *R*_2_, *R*_3_) − *R*_*A*_) + 0.014 * *R* − 0.0059 * *R*^2^ (dashed lines). **(E)** Deviation between simulated mean position of accessible volume (FPS software) and estimated mean position (SSM approach) including the correction factor *ε* (mean value: red line). **(F)** Distance between corrected screening mean position (SSM approach) and simulated mean position (FPS software) results in a deviation of 2.7 Å (RMSD). The large deviations for some positions (>6 Å) result from failed FPS simulations (unreasonable small accessible volumes due to interfering atoms close to the linker attachment site).

**Supplementary Figure S8.**
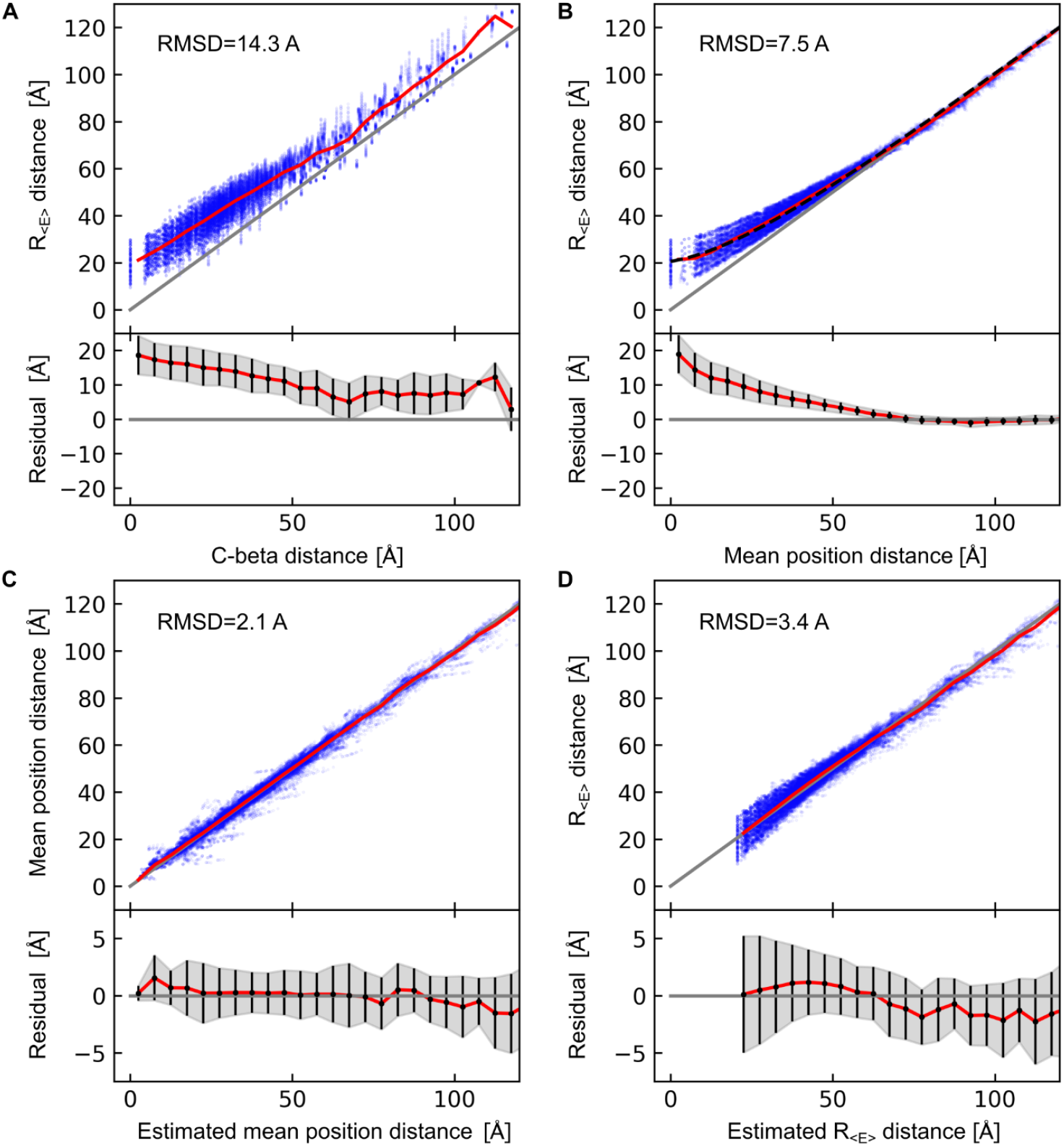
Correction parameter for distance simulation. **(A)** C-beta distances are plotted against simulated distances with FPS-software (blue datapoints) with mean values (red line). The bottom axis shows the mean residual (red line) and the standard deviation interval (error bars / gray area) on binned data from the top. **(B)** Mean dye position distances *R*_*M*P_ (center of mass of the AV-simulation) are plotted against the FRET-averaged distances (blue datapoints) with mean values (red line). The mean values are fitted to the curve 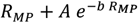 with *A* = 20.6 Å and *b* = 0.037 1/Å (black dashed line). **(C)** Mean dye position estimations based on the spherical sector calculation (SSM) approach are plotted against mean dye position distances *R*_*M*P_ from FPS-simulation software. **(D)** Mean dye position distances from the SSM-estimation are converted to FRET-averaged distances with the correction factors from (B) and plotted against the simulated 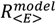 from FPS-simulation.

**Supplementary Figure S9.**
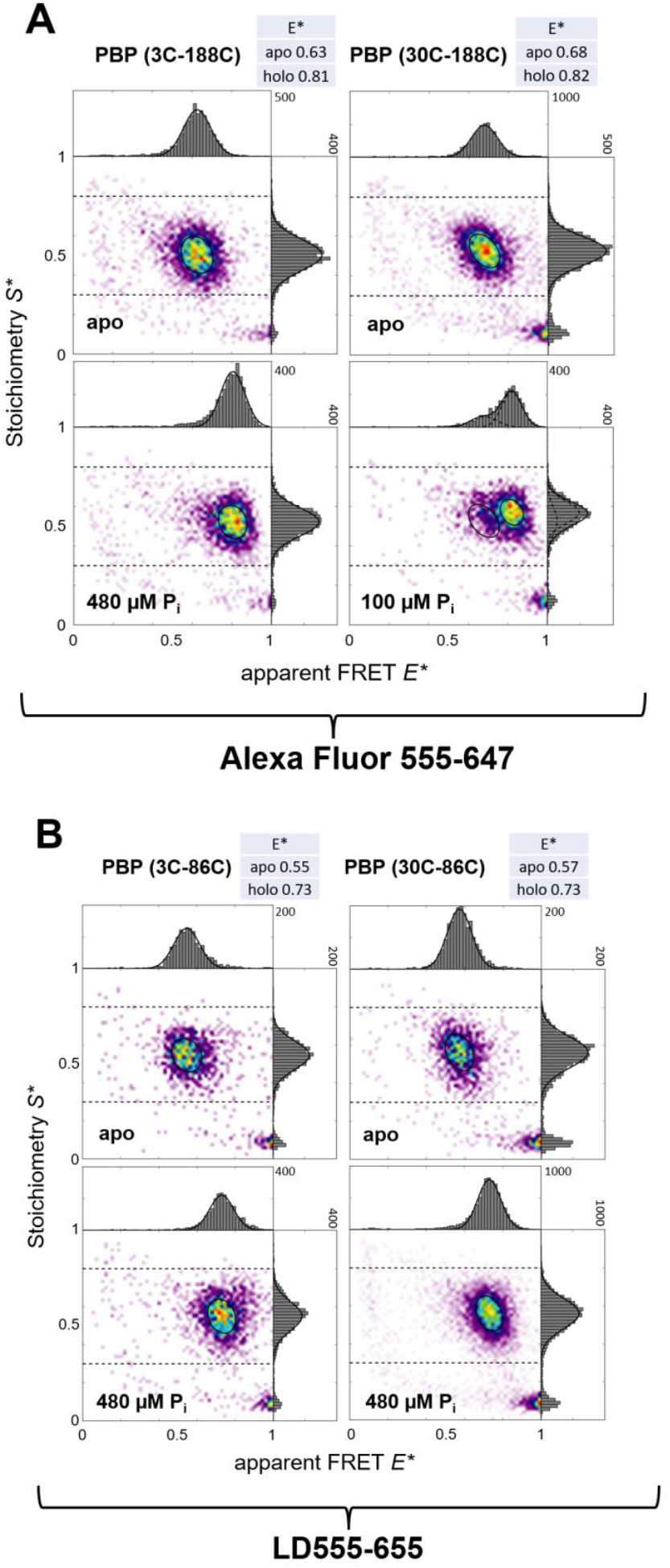
smFRET Characterization of four distinct PBP double-cysteine variants with two different fluorophore pairs with similar Förster radius. ALEX histograms with 61 bins of apo and holo states of PBP variants as indicated labelled with **(A)** Alexa Fluor 555-647 and **(B)** LD555-655 dyes. Mean values for E* are background corrected apparent FRET efficiencies analyzed by a dual-colour burst search with additional per-bin thresholds of all photons >150.

**Supplementary Figure S10.**
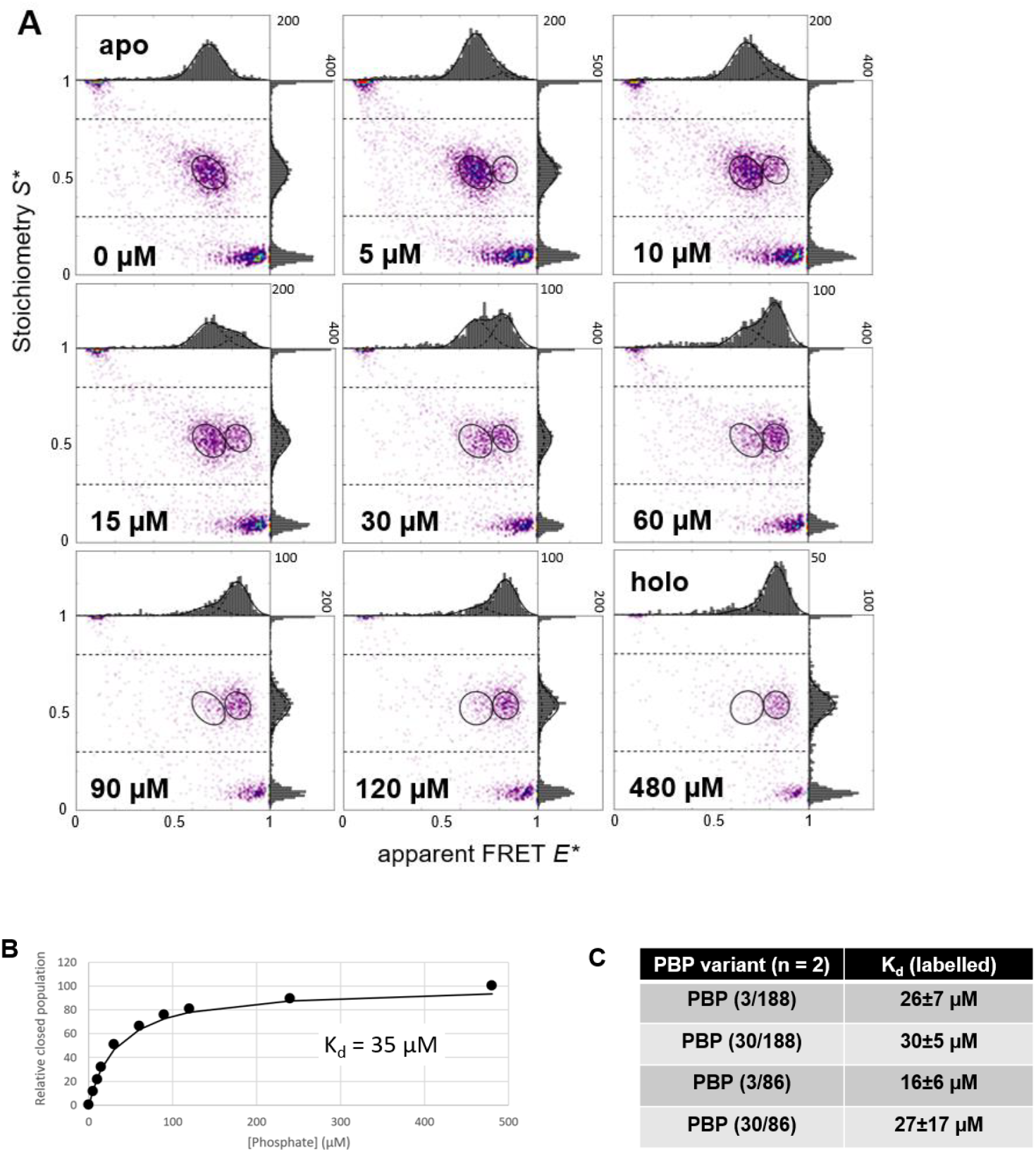
Biochemical characterization of labelled PBP variants using smFRET. **(A)** Representative data of S3-188C labelled with Alexa Fluor 555-647 at indicated phosphate concentrations including a two-state fit of low FRET apo and high-FRET holo state. **(B)** The binding curve was calculated from smFRET measurements considering the closed fraction (r_closed_) as a function of the substrate concentration. **(C)** Determined mean Kd-values for all four labelled PBP variants including standard deviation.

**Supplementary Figure S11.**
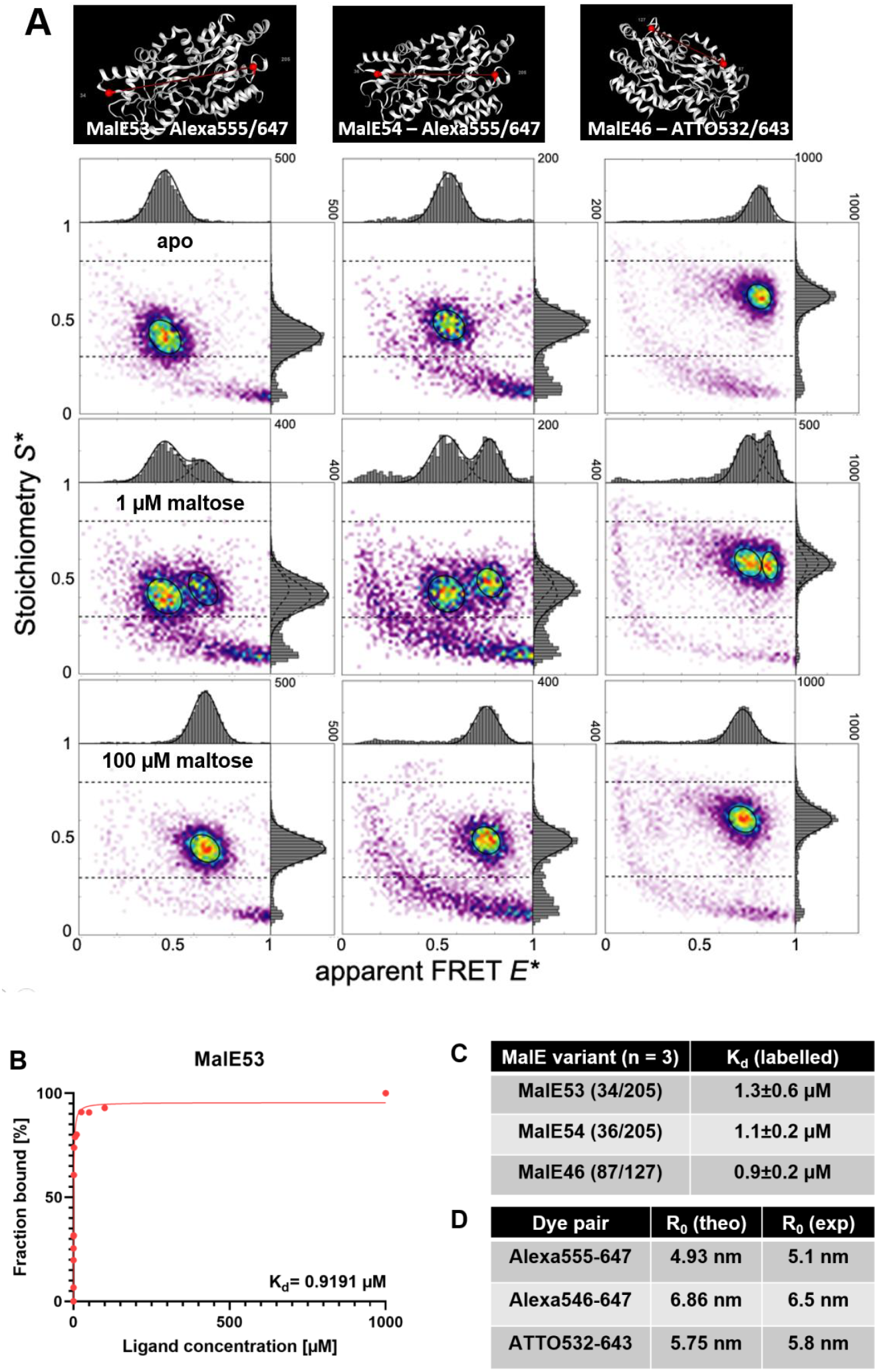
New smFRET and biochemical data of MalE variants used in accurate FRET analysis. **(A)** Label positions and smFRET of the respective variant for ligand-free apo, 1 µM maltose and saturated maltose (100 µM, holo). **(B)** Representative results from affinity titrations of MalE 53 with Alexa555-647 and **(C)** mean and standard deviation of determined Kd-values for all three labelled variants.

**Supplementary Figure S12.**
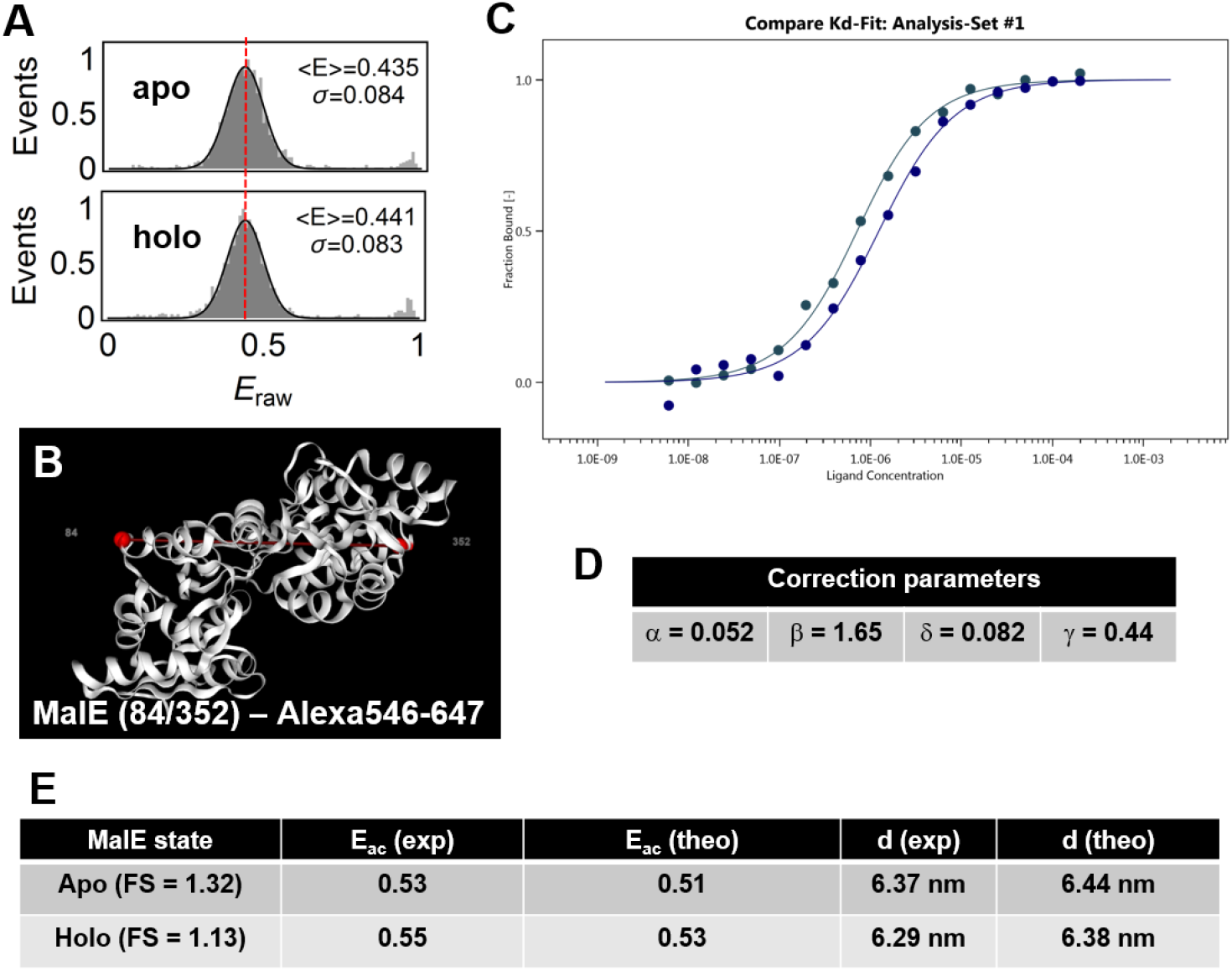
Biophysical and biochemical characterization of MalE variant with optimized FRET efficiency values but minimal FRET changes between apo and holo state. **(A)** Unccorrected FRET efficiency histogram of **(B)** MalE (84/352) labeled with Alexa Fluor 546-647 where ligand binding was verified by label-free microscale thermophoresis **(C)**. Conversion of the data into accurate FRET efficiency values was done using additional data with distinct FRET efficiency values (yet use of the same fluorophore pair on MalE) and resulted in correction parameters **(D)** and accurate FRET efficiency values and distances using a Förster radius of 6.5 nm **(E)**.

### D) Supplementary Tables 7-9

**Supplementary Table S7.**
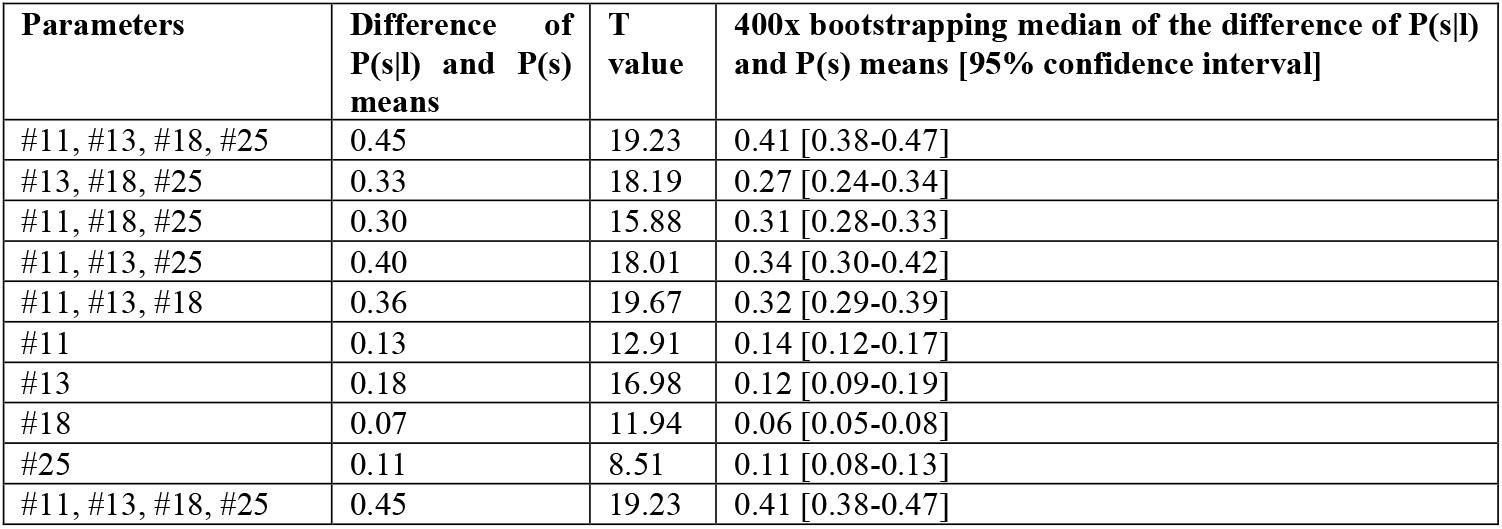
Parameter analysis. We compared the baseline model, the model with one of the four parameters removed and the predictive power of each parameter on its own. We show the difference between the means and the T value when comparing the distributions of P(s) and P(s|l), the average difference between the means after 400x bootstrapping together with the 95% CI intervals. The difference of means in our bayesian model is a metric for the model’s ability to separate the labeled residues from the background. Note that the median derived from bootstrapping of the differences of the distribution means diverges from the main analysis.

**Supplementary Table S8.**
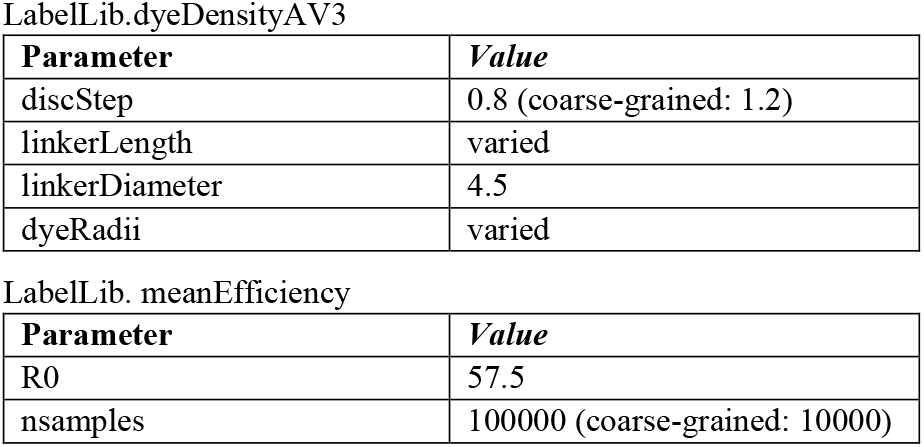
FPS settings. Overview of all user parameters set for the runtime analysis of the distance refinement simulation with the FPS software^16^. The default settings for the labelizer package and webserver use discStep = 0.8 and nsamples = 100000 (all other parameters depend on the selected fluorophore pair).

| Parameter | Value |
| --- | --- |
| discStep | 0.8 (coarse-grained: 1.2) |
| linkerLength | varied |
| linkerDiameter | 4.5 |
| dyeRadii | varied |

LabelLib.meanEfficiency
| Parameter | Value |
| --- | --- |
| R0 | 57.5 |
| nsamples | 100000 (coarse-grained: 10000) |

**Supplementary Table S9.** Spherical sector vs. FPS runtime comparison. Calculation time overview of the fast screening method (spherical sector calculation) and a coarse-grained distance refinement simulation (FPS software^16^) with 3500 distance pairs per pdb-file (100 distances, 35 different dye parameter) and a refined FPS simulation with 1400 distance pairs per pdb-file (40 distances, 35 different dye parameter).

| PDB | Molar mass [u] | SSM time per DA-pair [ms] | Coarse-grained FPS time per DA-pair [ms] | FPS time per DA-pair [ms] |
| --- | --- | --- | --- | --- |
| 3L6G | 28,830 | 0.81±0.04 | 103±5 | 315±11 |
| 2KHO | 65,650 | 0.84±0.01 | 166±5 | 419±8 |
| 2CG9 | 188,730 | 0.94±0.01 | 303±12 | 647±10 |
| 4B1O | 831,160 | 4.31±0.09 | 1113±36 | 2087±46 |
| 172L | 18,730 | 0.67±0.01 | 77±2 | 249±10 |
| 2A65 | 59,750 | 0.93±0.04 | 169±5 | 474±9 |
| 1WDN | 25,130 | 0.78±0.01 | 100±3 | 315±4 |
| 5XPD | 33,310 | 0.67±0.02 | 94±3 | 282±10 |
| 1P7B | 74,510 | 0.80±0.01 | 156±12 | 398±14 |
| 1HKA | 17,970 | 0.81±0.04 | 81±2 | 259±11 |

## Footnotes

1 Note that we also included some spin labels or biotin-linked fluorophores, yet these represent <5% of all labels in the database (see Supplementary Figure S1).

2 Since we deal with categorical data (e. g. secondary structure) and numerical data (e. g. relative surface area), we used Pearson correlation, interclass correlation and Cramer’s V for the combinations of numeric-numeric, categorical-numeric, categorical-categorical values, respectively (see methods for details).

3 We note here that it would be beneficial to compare the label scores of successfully labeled residues with non-successfully labeled residues in the future. However, we do not have the information on non-successfully labeled residues and only 1% of the considered residues (396 out of 43357) are known as labeled, which should not affect the comparison significantly.

4 An overview of experimentally determined and theoretical Förster radii is provided in Supplementary Figure S11D.

5 Additional notes were gathered to account for issues such as: (i) dimer and polymer protein structures, which were crystallization artefacts and needed to be deleted for structural analysis; (ii) missing residues in protein structure, i.e., when parts of the protein were not resolved completely; (iii) we identified inconsistencies or missing information

